# Characterization of Sec14 domain–containing proteins in the malaria parasite *Plasmodium falciparum*

**DOI:** 10.64898/2026.07.07.736992

**Authors:** Florian Lauruol, Dominik Šťastný, J. Pedro Fernandez-Murray, Christopher R. McMaster, Peter Griač, Dave Richard

**Affiliations:** Department of Microbiology-Infectious Diseases and Immunology, Faculty of Medicine, Université Laval, Québec, QC, Canada; CHU de Québec-Université Laval Research Centre, 2705 Boul. Laurier, Québec, QC, G1V 4G2, Canada; Centre de Recherche en Infectiologie de l’Université Laval, Quebec City, QC, Canada; Institute of Animal Biochemistry and Genetics, Centre of Biosciences, Slovak Academy of Sciences, Dúbravská cesta 9, 840 05, Bratislava, Slovakia; Department of Pharmacology, Dalhousie University, Halifax, NS, Canada

**Keywords:** malaria, plasmodium, phosphoinositides, sec14, PITP

## Abstract

Malaria, of which the most virulent form is caused by *Plasmodium falciparum* parasites, remains a major global health burden. The appearance of resistance to first line treatments artemisinin-based therapies, emphasizes the need to identify new parasite vulnerabilities to develop new therapeutics. Phosphoinositides are central regulators of membrane identity, vesicular trafficking, and signaling, and their synthesis depends on tightly controlled phosphatidylinositol transfer by Sec14-like phosphatidylinositol transfer proteins in many eukaryotes, yet their roles in *P. falciparum* remain poorly defined. Here, we analyzed six *P. falciparum* Sec14 domain–containing proteins: PfSec14-1 (PF3D7_0626400), PfSec14-2 (PF3D7_0629900), PfSec14-3 (PF3D7_0717100), PfSec14-4 (PF3D7_0920700), PfSec14-5 (PF3D7_1007200), and PfSec14-6 (PF3D7_1127600). Domain organization segregates these proteins into a BNIP-2 and Cdc42GAP homology (BCH) subfamily (PfSec14-3, PfSec14-5) and a canonical Sec14 subfamily (PfSec14-1, PfSec14-2, PfSec14-4, PfSec14-6). Yeast complementation assays showed that PfSec14-1, PfSec14-4, and PfSec14-6 partially rescue growth of a temperature-sensitive sec14 mutant, suggesting phosphatidylinositol and phosphatidylcholine transfer activity. Gene disruption revealed that PfSec14-1 is important for asexual blood-stage proliferation, whereas PfSec14-2 is dispensable under standard culture conditions. In contrast, mislocalization of PfSec14-1 and PfSec14-4 using a knock-sideways approach did not impair asexual growth. Subcellular localization indicates distinct distributions for PfSec14-1, PfSec14-2, and PfSec14-4. Together, these findings reveal functional and spatial diversification of Sec14-like phosphatidylinositol transfer proteins in *P. falciparum*.

## Introduction

Despite substantial reductions in global malaria mortality and morbidity over the past decade, malaria remains a major health challenge, with an estimated 610,000 deaths out of 282 million cases in 2024, mostly attributable to *Plasmodium falciparum*, the species responsible for the most severe clinical presentations (World Health Organization 2025). The widespread emergence of resistance to frontline antimalarial therapies, particularly artemisinin-based treatments, further underscores the urgent need for new therapeutic strategies (Rosenthal, Asua, and Conrad 2024).

To invade and replicate within host erythrocytes, *Plasmodium* parasites rely on tightly regulated processes of membrane biogenesis, intracellular signaling, vesicular trafficking, and hemoglobin digestion, all of which are fundamentally dependent upon lipid metabolism (Gulati et al. 2015; Tokumasu et al. 2021; Shunmugam et al. 2022). Among membrane lipids, phosphatidylinositol (PI) is rare but critically important as the precursor for phosphoinositides (PIPs), a versatile family of seven signaling lipids produced by phosphorylation at distinct positions on the inositol ring (Casares, Escribá, and Rosselló 2019). PIPs are central regulators of membrane identity, trafficking, cytoskeletal dynamics, and cell signaling (Hammond and Burke 2020). Key examples include PI4P, which is generated at the Golgi apparatus and the plasma membrane (PM) by PI4-kinases (PI4Ks) through the phosphorylation of PI. Interestingly, the intrinsic catalytic activity of PI4Ks is often insufficient to maintain the physiological pools of PI4P required for cellular function, a limitation attributed to low local concentrations of PI substrate or suboptimal catalytic efficiency (Zewe et al. 2020; Lete et al. 2020).

To overcome this limitation, cells utilize phosphatidylinositol transfer proteins (PITPs), members of the conserved lipid transfer protein (LTP) superfamily, which transfer PI to PI4Ks to stimulate synthesis and maintain a biologically sufficient PI4P pool (Vytas A. Bankaitis, Mousley, and Schaaf 2010; Kim et al. 2024). The Sec14-like and START-like families represent the two major PITP subclasses, with Sec14 proteins best known for integrating PIP signaling with vesicle formation and membrane homeostasis (Grabon, Bankaitis, and McDermott 2019). The *Saccharomyces cerevisiae* Sec14 protein (*Sc*Sec14p) is the founding and best-studied member of the Sec14-like PITP family. *Sc*Sec14p has been extensively characterized thanks to the discovery of the *S. cerevisiae sec14^ts^* strain, which harbors the temperature-sensitive *sec14-1* allele (V. A. Bankaitis et al. 1989). When grown at non-permissive temperatures above 34°C, *Sc*Sec14p function is inactivated and, because this protein is essential for the yeast, the strain is unable to grow. Members of the Sec14-like PITP family feature large hydrophobic pockets capable of binding PI or a second lipid (such as phosphatidylcholine (PC), phosphatidylserine (PS), or cholesterol) in a mutually exclusive manner (Schaaf et al. 2008; Ren et al. 2014). Another major group of LTPs involved in PIP metabolism is the oxysterol-binding protein (OSBP)-related protein (ORP) family, which contains an OSBP-related domain. ORPs are conserved across eukaryotes and participate in lipid exchange at membrane contact sites, mediating the transfer of sterols, PIPs, and phospholipids through PI4P-dependent mechanisms (Raychaudhuri and Prinz 2010; Lipp et al. 2020; Arora, Taskinen, and Olkkonen 2022). Beyond lipid transport, they play regulatory roles in cell signaling, proliferation, and homeostasis (Lin et al. 2023). In *S. cerevisiae*, seven ORP homologs, Osh1p–Osh7p, are critical for maintaining lipid balance and vesicular trafficking. Although the loss of a single *OSH* gene is tolerated, simultaneous deletion of all seven is lethal due to severe disruption of sterol homeostasis (Beh et al. 2001; Raychaudhuri et al. 2006; Sullivan et al. 2006). In *P. falciparum*, a single ORP-containing homolog (PF3D7_1131800) has been identified, though gene disruption studies indicate it is dispensable during the intraerythrocytic cycle (Mukherjee et al. 2022).

As in other eukaryotes, PIPs are critical in a number of essential cellular processes in *Plasmodium*. Pharmacological inhibition of PI4K has been shown to disrupt parasite growth, reduce gametocyte maturation, and impair transmission, making it an attractive antimalarial target (McNamara et al. 2013; Li et al. 2024). Phosphoinositide 3-kinase (PI3K) is essential for parasite survival and PI3P is linked to haemoglobin trafficking, resistance to artemisinin and apicoplast inheritance (Tawk et al. 2010; Vaid et al. 2010; Lu et al. 2020; Al Monla et al. 2025). Because PITPs are widely conserved in eukaryotes with evidence of conserved molecular functions from yeast to mammals (Hsuan and Cockcroft 2001; Phillips et al. 2006; Wyckoff, Solidar, and Yoder 2010; Grabon, Bankaitis, and McDermott 2019), it is likely that they are involved in PIP metabolism in *Plasmodium*. Interestingly, small molecule inhibitors (SMIs) targeting Sec14 PITPs are effective against different fungal pathogens in both *in vitro* and *in vivo* models. The major advantage of these SMIs is their high specificity, as they seem able to target a particular Sec14 protein selectively without influencing others (Nile et al. 2014; Chen et al. 2023; F. Zhang et al. 2020). Studying the role of PITPs in *Plasmodium* would provide a better understanding of PIP metabolism in this parasite and may unveil novel therapeutic avenues.

Not much is known with regards to the roles of Sec14-like PITPs in *Plasmodium* parasites. In one study, inhibition of the cyclic guanosine monophosphate (cGMP)-dependent protein kinase (PKG), a key regulator of parasite egress, motility, and Ca² signaling, was used to assess its impact on phosphorylation events in *P. berghei* ookinetes (Brochet et al. 2014). Phosphoproteome analysis revealed that a Sec14 protein encoded by PBANKA_1125200 (the ortholog of *P. falciparum* PF3D7_0626400 encoding PfSec14-1) exhibited reduced phosphorylation when PKG activity was blocked. This finding suggests that PBANKA_1125200 may be regulated by PKG-dependent phosphorylation, which could in turn modulate PI4P production and, indirectly, the levels of other PIPs. More recently, the biochemical properties of two *P. falciparum* Sec14 domain-containing proteins were characterized (Šťastný et al. 2025). PfSec14-2, encoded by PF3D7_0629900 displayed lipid-transfer activity for cholesterol, PI(4,5)P, and PC, and functionally complemented the absence of all seven yeast Osh proteins, suggesting a sterol–PIP exchange function. In contrast, PfSec14-6, encoded by PF3D7_1127600 transferred PS and complemented the *sec14^ts^* yeast strain, indicating activity more akin to the canonical yeast Sec14p protein (PI/PC transfer).

In this study, we conducted an initial exploration of the six *P. falciparum* Sec14 domain–containing proteins (PfSec14-1 encoded by PF3D7_0626400; PfSec14-2, PF3D7_0629900; PfSec14-3, PF3D7_0717100; PfSec14-4, PF3D7_0920700; PfSec14-5, PF3D7_1007200; and PfSec14-6, PF3D7_1127600). We show that these proteins can be separated into two groups based on the spatial organization of their Sec14 domains: *Pf*Sec14-3 and *Pf*Sec14-5 belong to the BNIP-2 and Cdc42GAP Homology (BCH) subfamily, while the other four belong to the canonical Sec14 family. Functional assays revealed that PfSec14-1, PfSec14-4, and PfSec14-6 can partially complement the absence of a functional *Sc*Sec14p in the *sec14^ts^*yeast strain, suggesting a PI/PC transfer activity for these proteins. This confirms what was already published for PfSec14-2 and PfSec14-6 (Šťastný et al. 2025). The disruption of PfSec14-1 led to a major growth defect in asexual stage parasites, whereas the PfSec14-2 protein was dispensable. Functional inactivation by mis-targeting PfSec14-2 and PfSec14-4 using the knock-sideways (KS) method did not have a measurable effect on asexual-stage parasite growth. Finally, subcellular localization demonstrated that PfSec14-1 exhibits dynamic nuclear and ER-Golgi localization, PfSec14-2 potentially associates with the PM, inner membrane complex (IMC), and basal complex (BC), and PfSec14-4 localizes to the ER and Golgi.

## Results

### *In silico* characterization of Sec14 domain-containing proteins in *P. falciparum*

As previously described, the *P. falciparum* genome encodes six proteins containing a Sec14 domain (Lauruol and Richard 2024), designated PfSec14-1 to PfSec14-6 (Table 1 and Fig. 1A). Canonical Sec14 proteins contain two domains: a Sec14 domain that forms the lipid binding pocket and a four-helix subdomain referred as the N-terminal Sec14 domain that plays a role in membrane interactions (Miller et al. 2015). Among the six Sec14 domain-containing proteins of the parasite, PfSec14-1, PfSec14-2, PfSec14-4, and PfSec14-6 contain exclusively the Sec14 domain and the N-terminal Sec14 domain (Lauruol and Richard 2024). In contrast, PfSec14-3 and PfSec14-5 lack the N-terminal Sec14 domain and each contain an additional module: PfSec14-3 harbors a Macro domain, while PfSec14-5 contains a Rho GTPase-activating protein (RhoGAP) domain (Lauruol and Richard 2024).

**Fig. 1.**
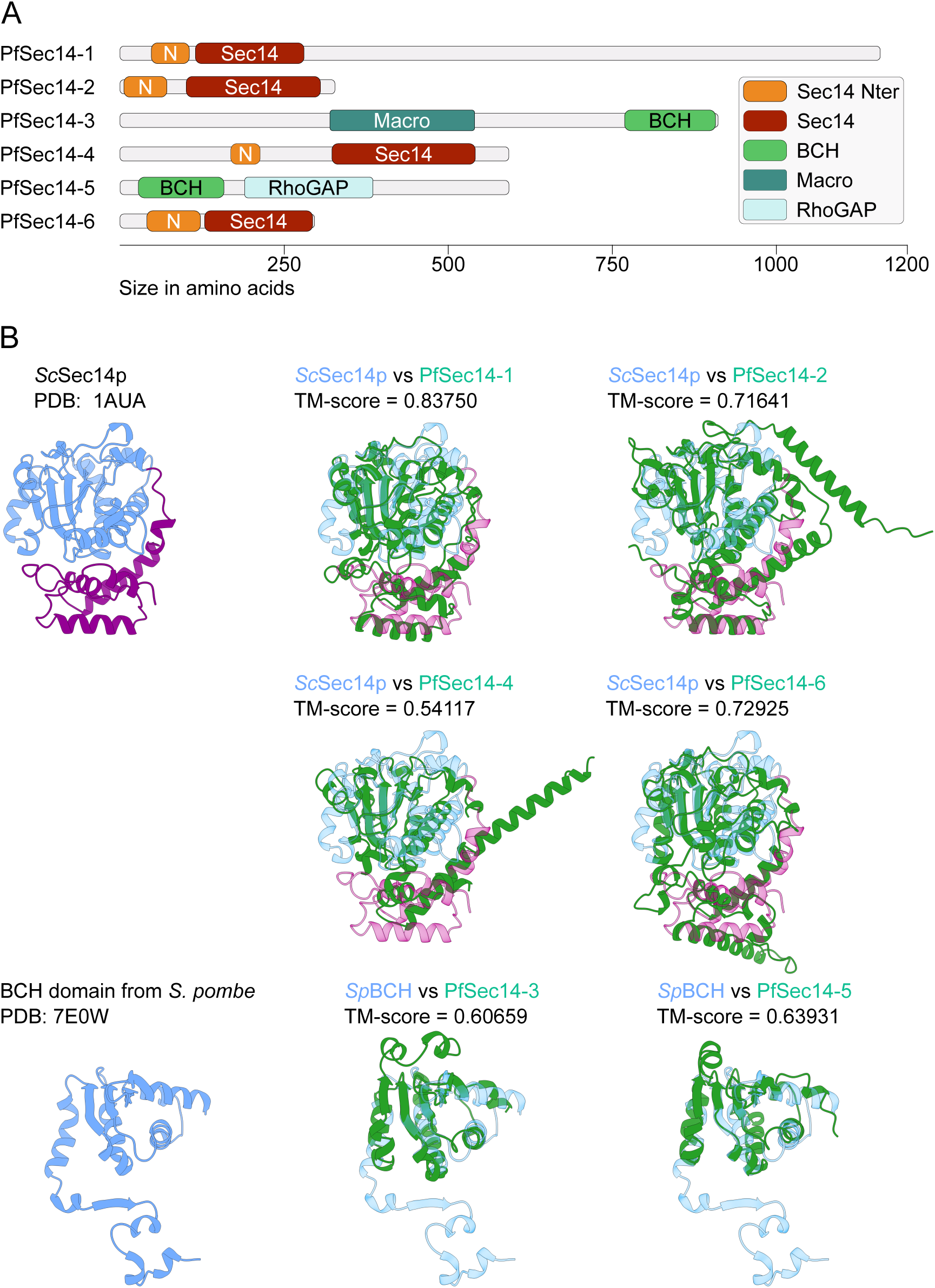
Structural predictions of Sec14 domain-containing proteins in *Plasmodium falciparum*. (A) Schematic representation of *P. falciparum* Sec14 proteins showing predicted domain organization. (B) AlphaFold3-predicted 3D structures of the Sec14 domains from PfSec14-1, PfSec14-2, PfSec14-4, and PfSec14-6 (green) superimposed on the crystal structure of the *S. cerevisiae* Sec14p (cyan: Sec14 domain, magenta: Sec14 N-terminal domain), and AlphaFold3-predicted BCH-like domains from PfSec14-3 and PfSec14-6 (green) superimposed on the BCH domain from *S. pombe* (cyan). N: Sec14 N-terminal domain. Sec14: Sec14 domain. Macro: Macro domain. BCH: BNIP-2 and Cdc42GAP Homology. RhoGAP: Rho GTPase activation protein domain. TM-score: template modeling score.

**Table 1.** List of *P. falciparum* Sec14 domain-containing proteins.

The divergence of PfSec14-3 and PfSec14-5, marked by the absence of canonical N-terminal Sec14 domains and the presence of Macro or RhoGAP domains, suggested they were part of the BCH subfamily of the Sec14 superfamily (Gupta et al. 2012). To validate this, Hidden Markov Model (HMM) profile searches were conducted against the *P. falciparum* Kyoto Encyclopedia of Genes and Genomes (KEGG) database, demonstrating closer relationships with BCH motifs than with canonical Sec14 domains (Table 2). BCH domains are functionally specialized for small GTPase regulation and associated signaling pathways, rather than phospholipid transfer activities (Chichili et al. 2021; Pan and Low 2012).

**Table 2.** Determination of whether Sec14 domains of PfSec14 proteins belong to the BCH or Sec14 family.

To assess structural conservation, structures were predicted for all *P. falciparum* Sec14 domain-containing proteins using AlphaFold 3 (Abramson et al. 2024) (Fig. S1). All predictions were robust with a predicted template modelling (pTM) score > 0.5 except for PfSec14-1 (pTM = 0.36), likely due to increased disorder within its large low-complexity regions (LCR). Nevertheless, the AlphaFold 3 algorithm predicts a Sec14 domain inside PfSec14-1 with strong confidence with all residues within this domain presenting a high predicted local distance difference test (pLDDT) score > 70. The Sec14/BCH domains were then extracted from these models and aligned to either the *S. cerevisiae* Sec14p structure (Sha et al. 1998) or the BCH domain of *Schizosaccharomyces pombe* (Chichili et al. 2021) using the US-align alignment tool (C. Zhang et al. 2022) (Fig. 1B). This showed that PfSec14-1, PfSec14-2, PfSec14-4 and PfSec14-6 Sec14 domains are homologous to *S. cerevisiae* Sec14p (TM-score > 0.5). By contrast, PfSec14-3 and PfSec14-5 structurally resemble the BCH domain of *S. pombe* (TM-score > 0.6).

This *in silico* characterization reveals substantial diversity within the *P. falciparum* Sec14 protein family. Comparative modeling demonstrates that PfSec14-1, PfSec14-2, PfSec14-4, and PfSec14-6 possess highly conserved Sec14 domains closely aligned with the canonical lipid-binding yeast Sec14p. In contrast, PfSec14-3 and PfSec14-5 exhibit greater structural resemblance to BCH-type domains, which are typically implicated in distinct signaling functions across other eukaryotic systems.

### Heterologous expression of *P. falciparum* Sec14 homologues can complements distinct lipid transfer proteins in *S. cerevisiae* mutants

To investigate their potential lipid transfer activity, we expressed all six *P. falciparum* Sec14 homologues in *S. cerevisiae* mutant strains to assess their ability to complement known phenotypes of yeast strains with defective LTPs of the Sec14p and Osh family. Due to gene size and sequence complexity, only the predicted Sec14 domains sequence of PfSEC14-1 and PfSEC14-4 were used while the full-length (FL) sequence was used for the 4 other genes. All proteins were C-terminally tagged with a Myc epitope to monitor expression. Western blot analysis confirmed the expression of all constructs at the expected molecular weights (Fig. S2). The predicted sizes for the Myc-tagged proteins were: PfSec14-1(dom): 35.8 kDa; PfSec14-2(FL): 39.3 kDa; PfSec14-3(dom): 21.4 kDa; PfSec14-4(FL): 71.8 kDa; PfSec14-5(FL): 68.6 kDa; and PfSec14-6(FL): 36.2 kDa. The observed bands on the blot matched these predictions, appearing at approximately 35, 40, 20, 70, 70, and 35 kDa, respectively (Fig. S2). Interestingly, the constructs expressing only the isolated Sec14 domains (PfSec14-1 and PfSec14-3) displayed significantly weaker signals compared to the full-length constructs of the other Sec14 proteins, suggesting potential instability or degradation of the domain-only constructs in yeast.

### PfSec14-1, PfSec14-4 and PfSec14-6 can partially complement the *sec14-1ts* growth defect

We first assessed the ability of *P. falciparum* Sec14 domain-containing proteins to rescue the growth defect of the temperature-sensitive *sec14-1^ts^* strain (Novick, Field, and Schekman 1980; Cleves et al. 1991). Yeast strains carrying the *sec14-1^ts^* allele do not grow at non-permissive temperatures above 34°C. At the permissive temperature (26°C), all transformants displayed normal growth. At the restrictive temperatures of 34°C and 37°C, the negative control (containing empty vector) failed to grow, while the positive control (expressing *Sc*Sec14) grew well (Fig. 2A). At the *sec14-1^ts^*non-permissive temperature of 34°C, expression of PfSec14-1 (Sec14 domain), PfSec14-4 (FL), and PfSec14-6 (FL) allowed for partial growth of the *sec14-1^ts^* strain (Fig. 2A). At 37°C, PfSec14-4 and PfSec14-6 retained the ability to support limited growth, whereas PfSec14-1 supported growth to a much lesser degree. It is interesting that despite the low protein levels detected for the PfSec14-1 domain (Fig. S2), this protein was still capable of partially rescuing the *sec14-1^ts^* growth defect. These results indicate partially overlapping functions between yeast Sec14p and these specific *P. falciparum* Sec14 domain-containing proteins. The absence of complementation by PfSec14-3 and PfSec14-5 reinforces the idea that these proteins likely possess a BCH domain not involved in phospholipid transfer activity.

**Fig. 2.**
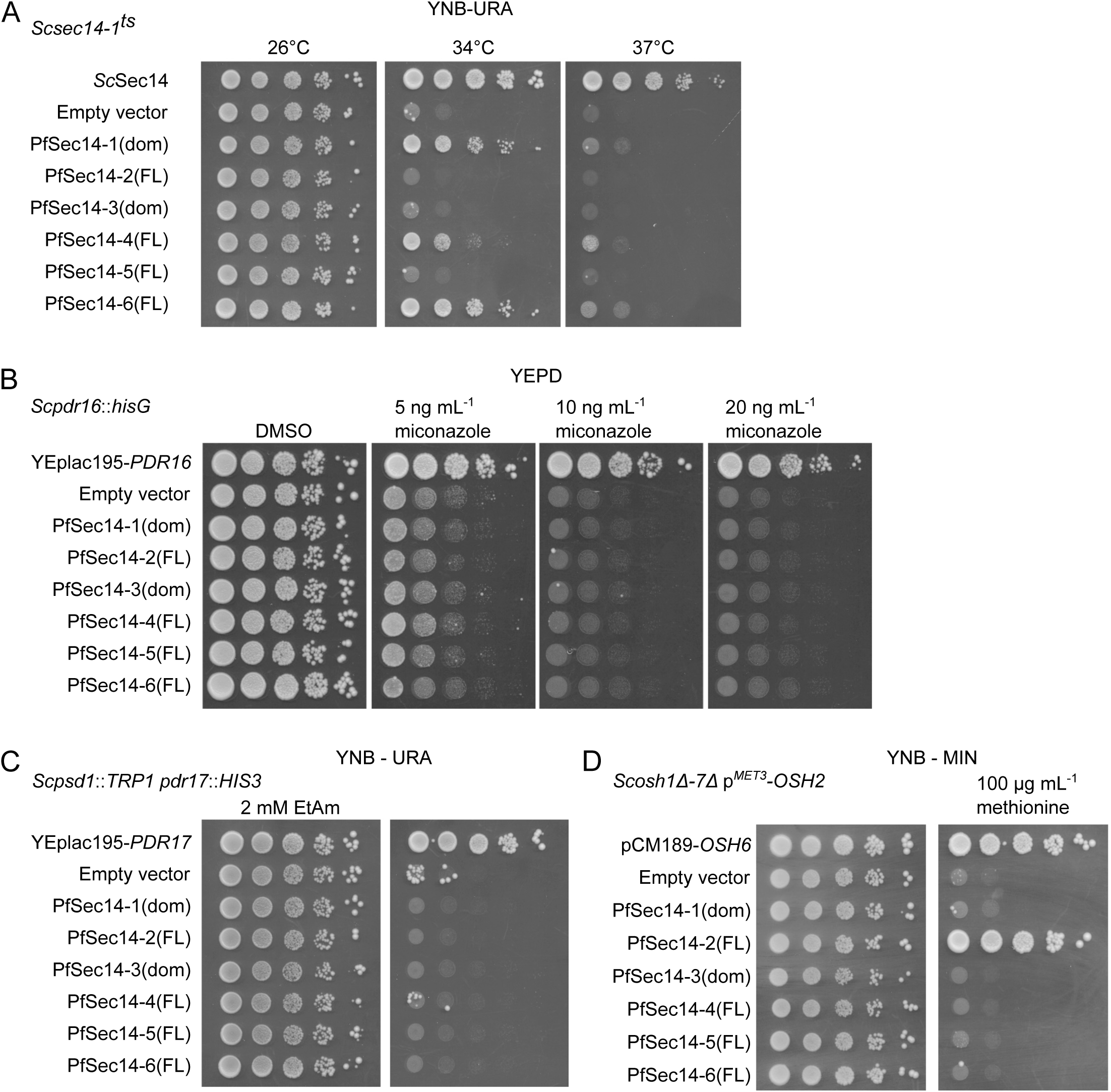
Complementation of *S. cerevisiae* lipid transfer protein mutant strains by overexpression of *P. falciparum* Sec14 homologues. (A) PfSec14-1, -4 and -6 partially rescue the growth defect of the *sec14-1^ts^* yeast strain at the non-permissive temperatures of 34°C and 37°C. (B) Expression of PfSec14 proteins does not rescue increased sensitivity of the *Scpdr16*::*hisG* strain to the imidazole antifungal miconazole. (C) Expression of PfSec14 proteins does not rescue the ethanolamine (EtAm) auxotrophy of the *Scpsd1*::*TRP1 pdr17*::*HIS3* strain, indicating absence of *pdr17Δ* complementation. (D) Only PfSec14-2 rescues the growth defect of the *Scosh1Δ-7Δ* strain.

*S. cerevisiae* possesses several homologs of Sec14p. Among these, Pdr16p (Sfh3p) is implicated in the yeast’s sensitivity to azole antifungals, whereas Pdr17p (Sfh4p) plays a role in phosphatidylethanolamine (PE) metabolism, making the absence of these proteins easily detectable through distinct phenotypes. Deletion of Pdr16p results in increased sensitivity to azole antifungals (Holič et al. 2014; van den Hazel et al. 1999), while deletion of Pdr17p causes a defect in the conversion of PS to PE, leading to ethanolamine auxotrophy in combination with the absence of functional PS decarboxylase Psd1p (Pevalová et al. 2019; Wu et al. 2000; Kannan, Riekhof, and Voelker 2015). To assess whether *P. falciparum* Sec14 domain–containing proteins could complement these phenotypes, we expressed them in *S. cerevisiae* strains lacking Pdr16p (*pdr16*::*hisG*) or Pdr17p (*psd1*::*TRP1 pdr17*::*HIS3*).

As expected, the *pdr16*::*hisG* strain exhibited increased susceptibility to the antifungal agent miconazole compared to the positive control (strain expressing yeast Pdr16p from the rescue plasmid YEplac195-*PDR16*). Expression of *P. falciparum* Sec14 domain–containing proteins did not alleviate this sensitivity, as *pdr16* transformants showed growth patterns comparable to the negative control (empty vector) in the presence of miconazole (Fig. 2B).

Similarly, absence of Pdr17p in the *psd1*::*TRP1 pdr17*::*HIS3* strain abolished growth in the absence of ethanolamine, in contrast to the positive control (*psd1*::*TRP1 pdr17*::*HIS3* strain expressing yeast Pdr17p from the rescue plasmid YEplac195-*PDR17*). None of the *P. falciparum* Sec14 domain–containing proteins restored growth under ethanolamine-depleted conditions, matching the phenotype of the negative control (empty vector). This result indicates that none of the *P. falciparum* Sec14 like proteins can functionally replace yeast Pdr17p (Fig. 2C).

### Only PfSec14-2 can complement *OSH* family deletion growth phenotype

Building on our previous finding that PfSec14-2 can rescue the loss of all Osh proteins in *S. cerevisiae* (Šťastný et al. 2025), we sought to determine whether any other *P. falciparum* Sec14 domain–containing proteins could similarly complement this essential function. To test this, we employed a yeast strain with all *OSH* genes deleted, containing an integrated *OSH2* gene under the control of the *MET3* promoter (Beh et al., 2001). When the *OSH2* gene is expressed in the absence of methionine, this strain is viable. Upon addition of methionine, which represses the *MET3* promoter, the strain fails to grow. The *osh1*Δ-*7*Δ p*^MET3^*-*OSH2* strain was transformed with the p416-GPD plasmids carrying genes encoding *P. falciparum* Sec14 domain-containing proteins. Only PfSec14-2 restored growth in the presence of methionine (Fig. 2D), indicating it can substitute for the collective essential function of all yeast Osh proteins.

In summary, these results demonstrate that some of the *P. falciparum* Sec14 proteins are capable of substituting for specific yeast lipid transfer activities connected to the action of Sec14p and Osh proteins. The lack of functional complementation of Sec14p observed in the *sec14-1^ts^* strain for both PfSec14-3 and PfSec14-5 is consistent with their Sec14 domains being closely related to the BCH domain and therefore likely lacking lipid-binding capability.

### *PfSEC14-1* gene disruption leads to severe growth defects in asexual stages

To investigate the function of PfSec14 proteins in *P. falciparum*, we performed targeted gene disruption (TGD) using the selection-linked integration (SLI) system (Birnbaum et al. 2017) (Fig. 3A). While PfSEC14-1 and PfSEC14-2 were successfully disrupted (Fig. 3B: both strains have proper 5’ and 3’integration events and absence of a WT allele, as detected by PCR), repeated attempts to disrupt PfSEC14-3, PfSEC14-4, PfSEC14-5, and PfSEC14-6 failed, as parasites did not recover from drug selection, suggesting that these genes may be essential. Interestingly, a whole genome PiggyBac insertion screen suggested that all Sec14 domain-containing genes except PfSEC14-3 are important for parasite asexual *in vitro* growth (Table 1) (M. Zhang et al. 2018). However, inactivation in *P. berghei* was successful for the PfSEC14-4 homolog (PBANKA_0821600), resulting in a slow-growth phenotype, and for the PfSEC14-5 homolog (PBANKA_1205400), with no apparent effect on growth. Inactivation of the PfSEC14-1 homolog (PBANKA_1125200) was not possible, suggesting an essential role for this gene in this parasite species (Table 1) (Bushell et al. 2017).

**Fig. 3.**
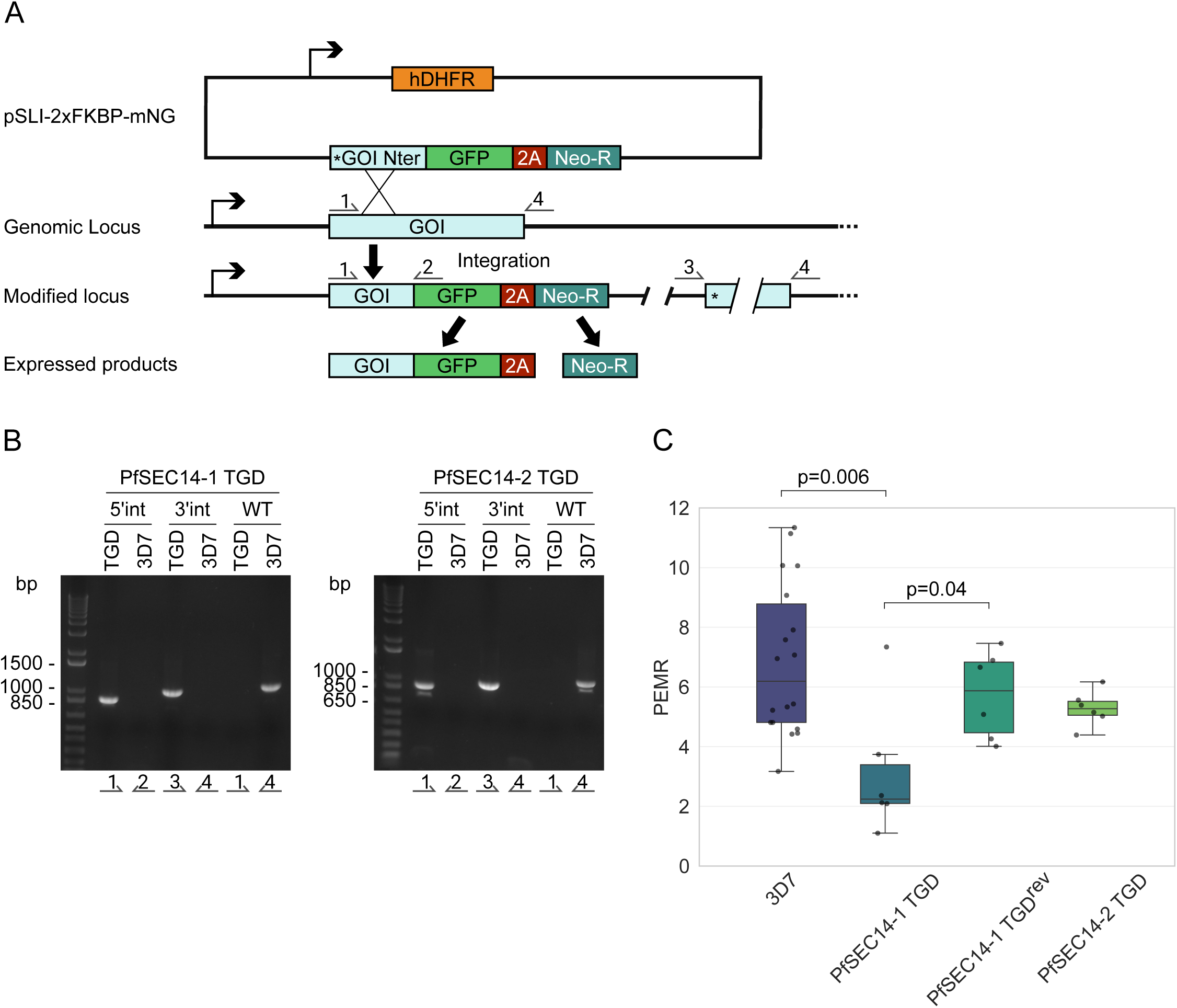
Disruption of the *PfSEC14-1* and *PfSEC14-2* genes. (A) Schematic of the disruption strategy by single cross-over recombination using SLI. Numbered arrows indicate the positions of primers used to verify integration. (B) PCR on parasite genomic DNA showing the proper integration of the TGD vector in the *PfSEC14-1* and *PfSEC14-2* loci and loss of the wild type locus. (C) Box plot showing the growth rate represented by the parasite erythrocyte multiplication rate (PEMR). Each condition was analyzed from three biological replicates. 3D7 mean PEMR ± SD = 6.86 ± 2.56. PfSEC14-1 TGD mean PEMR = 3.12 ± 2.23. PfSEC14-1^rev^ mean PEMR = 5.73 ± 1.47. PfSEC14-2 TGD mean PEMR = 5.28 ± 0.59. Only statistically significant differences between strains are indicated (Mann–Whitney U test). hDHFR: human dihydrofolate reductase. GOI: gene of interest. Nter: N terminal. GFP: green fluorescent protein. 2A: 2A peptide. Neo-R: neomycin resistance. *: stop codon. WT: wild type. TGD: targeted gene disruption. bp: base pair. PEMR: parasite erythrocyte multiplication rate

The growth of the PfSEC14-1 and PfSEC14-2 TGD parasite lines was monitored over 120 hours (corresponding to two and a half intraerythrocytic cycles) and compared with the parental 3D7 strain (Fig. 3C and Fig. S3). The PfSEC14-2 TGD line displayed no significant difference in parasite erythrocyte multiplication rate (PEMR) relative to 3D7 (PEMR ± SD = 5.28 ± 0.59 vs. 6.86 ± 2.56; Mann–Whitney U test, p = 0.3336, n = 3), indicating that PfSEC14-2 is dispensable for *in vitro* asexual blood-stage development, consistent with PiggyBac data showing successful transposon insertions within multiple regions of the gene coding sequences (CDS), although those insertions were associated with a strong negative fitness effect (mutant fitness score = −2.87) not reflected in our results (Table 1) (M. Zhang et al. 2018). In contrast, disruption of PfSEC14-1 resulted in a marked growth defect, with the PfSEC14-1 TGD strain exhibiting a PEMR approximately 50% lower than that of 3D7 (3.12 ± 2.23 vs. 6.86 ± 2.56; Mann–Whitney U test, p = 0.0057, n = 3). However, after several weeks in continuous culture, the PfSEC14-1 TGD parasites reverted to wild type-like growth, as indicated by comparable PEMR values between the revertant line (PfSEC14-1 TGD^rev^) and the 3D7 control (5.73 ± 1.47 vs. 6.86 ± 2.56; Mann–Whitney U test, p = 0.2712, n = 3).

Because the PfSec14-1 TGD line reverted to a wild type–like growth phenotype before cultures reached the densities required for invasion and attachment assays, and because multiple independent frozen aliquots failed to re-establish robust growth after thawing and gradually died, we were unable to expand cultures for these downstream experiments. However, within the limited window prior to reversion, we quantified the number of merozoites per mature schizont and detected no significant difference compared with the 3D7 parental line (Fig. S4A) (merozoites per schizont ± SD = 23.72 ± 2.90 vs. 24 ± 2.99; n = 35), and morphologically both schizonts and their merozoites appeared normal, indicating that reduced growth was not explained by gross defects in schizogony.

To try to understand why this TGD line reverted to a wild type–like growth phenotype, we used quantitative RT-PCR to assess whether this reversion could be explained by compensatory overexpression of the other *Sec14* genes (Fig. S4B). No expression of the *PfSEC14-1* gene was detected after reversion, indicating that the phenotype was not due to re-expression of the wild type gene or outgrowth of a wild type contaminant population. Nevertheless, no significant differences in transcript abundance for any of the other genes were observed relative to the 3D7 control. Notably, PfSEC14-2 and PfSEC14-4 transcript levels showed high variability among PfSEC14-1 TGD^rev^ replicates, indicating fluctuating expression without a consistent, statistically supported trend. At most, this pattern is compatible with, but does not demonstrate, a possible compensatory adjustment of these proteins in some replicates and therefore cannot be interpreted as firm evidence for transcriptional compensation.

Globally, this analysis suggests that PfSEC14-1 is important for parasite growth, as its disruption leads to a severe growth defect. In contrast, PfSEC14-2 disruption does not affect parasite growth, suggesting that this member is dispensable for asexual erythrocytic stage development. The failure to generate TGD for PfSEC14-3, PfSEC14-4, PfSEC14-5 and PfSEC14-6 indicates that these genes may play essential roles during in vitro asexual replication.

A limitation of the TGD approach to study PfSEC14-1 function is the instability of this line, which reverted to a wild type–like growth phenotype and frequently failed to recover after freezing. As a consequence, we could not perform invasion or attachment assays, and our functional conclusions are restricted to overall asexual growth rate and merozoite number per schizont. Instability of genetically modified *Plasmodium* lines over time in culture has been noted previously (Knuepfer et al. 2017) and underscores the importance of alternative conditional strategies.

### Induced mislocalization of PfSec14-2 and PfSec14-4 does not measurably impact parasite proliferation during the asexual intraerythrocytic cycle

Due to instability of the PfSEC14-1 TGD phenotype and the failure to obtain TGD lines for PfSEC14-3, PfSEC14-4, PfSEC14-5 and PfSEC14-6, we next pursued conditional approaches. Conditional KS enables the rapid mislocalization of FK506 binding protein (FKBP)-tagged proteins via rapalog-induced FKBP–FKBP12-rapamycin-binding (FRB) heterodimerization, thereby reducing access to their native site of action (Hughes and Waters 2017; Birnbaum et al. 2017) (Fig. 4A). We attempted C-terminal tagging of *PfSEC14* genes with 2xFKBP-mNeonGreen (mNG) and obtained 2xFKBP-mNG lines for *PfSEC14-1*, *PfSEC14-2*, and *PfSEC14-4*, whereas integrants could not be recovered for *PfSEC14-3*, *PfSEC14-5*, or *PfSEC14-6*. Following cloning by limiting dilution, PCR confirmed correct integration and loss of the WT locus for PfSEC14-2-2xFKBP-mNG and PfSEC14-4-2xFKBP-mNG lines (Fig. 4B). In contrast, PfSEC14-1-2xFKBP-mNG retained a detectable WT locus, which prevented KS-based phenotyping but still allowed localization analysis via the mNG tag (see below). To try to obtain a fully modified conditional line for PfSEC14-1, we generated a conditional knockdown (KD) parasite line (PfSEC14-1^TetR^) using the TetR-aptamer system, which enables tetracycline-dependent translational control (Ganesan et al. 2016; Rajaram, Liu, and Prigge 2020) (Fig. S5A). PCR confirmed correct integration of the KD construct and the absence of the WT locus (Fig. S5B).

**Fig. 4.**
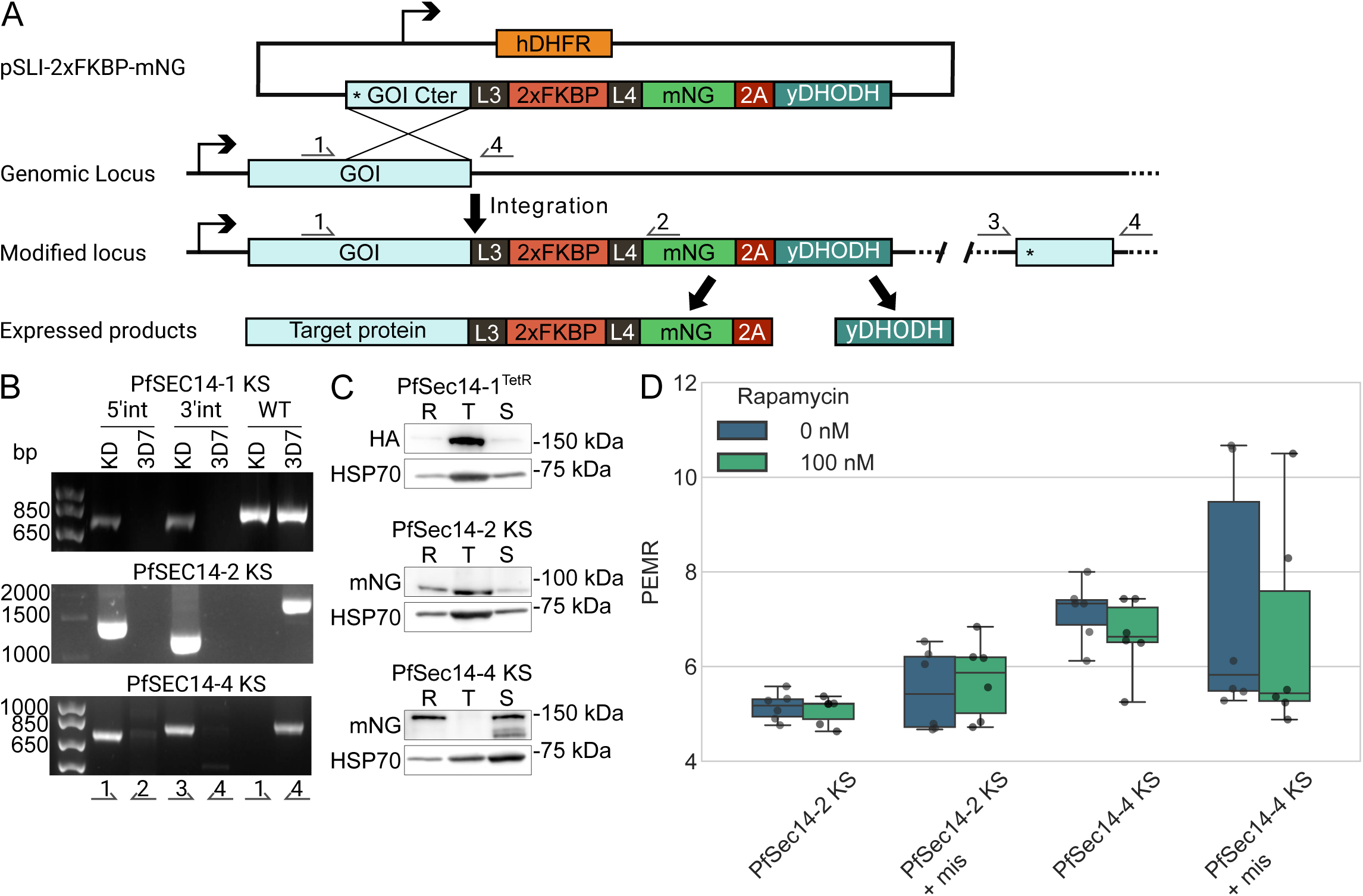
Generation of PfSec14-1, PfSec14-2, and PfSec14-4 knock-sideways lines. (A) Illustration of the single crossover recombination strategy employed for gene tagging using the SLI system. Numbered arrows indicate the positions of primers used to verify integration. (B) PCR-based validation of tagging vector integration at the *PfSEC14-1*, *PfSEC14-2* or *PfSEC14-4* loci performed on parasite genomic DNA. (C) Western blot of protein extracts from ring, trophozoite, and schizont stages. PfSec14-1 expression in the PfSec14-1^TetR^ line was detected at the expected size of approximately 140 kDa with an anti-HA antibody. An anti-mNG antibody was used to detect PfSec14-2 and PfSec14-4 in KS lines, showing bands at the expected sizes of approximately 90 kDa for PfSec14-2 and 120 kDa for PfSec14-4. (D) Box plot showing growth rate as parasite erythrocyte multiplication rate (PEMR). Each condition was analyzed from three biological replicates. PfSec14-2 KS mean PEMR = 5.15 ± 0.30 (0 nM rapamycin) and 5.07 ± 0.29 (100 nM rapamycin). PfSec14-2 KS + mis mean PEMR = 5.50 ± 0.87 (0 nM rapamycin) and 5.72 ± 0.84 (100 nM rapamycin). PfSec14-4 KS mean PEMR = 7.16 ± 0.65 (0 nM rapamycin) and 6.65 ± 0.80 (100 nM rapamycin). PfSec14-4 KS + mis mean PEMR = 7.28 ± 2.61 (0 nM rapamycin) and 6.63 ± 2.26 (100 nM rapamycin). No statistically significant difference was observed between non induced (0 nM) and induced (100 nM) KS conditions (Mann–Whitney U test). hDHFR: human dihydrofolate reductase. yDHODH: yeast dihydroorotate dehydrogenase. mNG: mNeonGreen. FKBP: FK506 binding proteins. L3/L4: linker. GOI: gene of interest. Cter: C terminal. 2A: 2A peptide. *: stop codon. WT: wild type. KD: knock down. bp: base pair. R: rings, T: trophozoites, S: schizonts. PEMR: parasite erythrocyte multiplication rate. mis: mislocalizer.

Expression of the tagged proteins was assessed by Western blot across ring, trophozoite, and schizont stages (Fig. 4C and Appendix A). PfSec14-1-2xFKBP-mNG was not detectable at the expected size (∼184 kDa) using anti-mNG (Appendix A). In contrast, in the PfSEC14-1^TetR^ line, PfSEC14-1-HA was readily detected at the expected size (∼140 kDa) across the intraerythrocytic cycle using anti-HA antibody (Fig. 4C and Appendix A). This discrepancy likely reflects low abundance of the tagged PfSec14-1-mNG, which prevents its detection with the anti-mNG antibody used, as this antibody appears substantially less sensitive (requiring much longer exposure to visualize a specific signal) than the anti-HA antibody. Transcriptomic (Kucharski et al. 2020; Chappell et al. 2020) and proteomic (Pease et al. 2013) datasets indicate that PfSEC14-1 is expressed throughout the intraerythrocytic cycle, with upregulation toward late stages and high protein abundance particularly in ring stages. Our Western blot, which is not quantitative, nonetheless shows the presence of the protein at all intraerythrocytic stages, in agreement with these transcriptomic and proteomic data. PfSec14-2-2xFKBP-mNG was detected at the expected size (∼90 kDa) throughout the intraerythrocytic cycle. This pattern is consistent with RNA-seq data (Kucharski et al. 2020; Chappell et al. 2020) showing substantial expression, with a peak at early stages followed by a decrease that remains relatively stable until the end of the cycle. Proteomic data likewise support the presence of PfSec14-2 in ring, trophozoite, and schizont stages, with slightly higher abundance in rings than in trophozoites and schizonts (Pease et al. 2013). PfSec14-4-2xFKBP-mNG was observed at the expected size (∼120 kDa) in rings and schizonts but not in trophozoites, and in schizonts an additional lower molecular weight band (∼100 kDa) was present, reflecting a possible stage-specific processing. This pattern aligns with transcriptomic (Kucharski et al. 2020; Chappell et al. 2020) and proteomic (Pease et al. 2013) data indicating higher expression of PfSEC14-4 in rings and schizonts compared with trophozoites.

To assess the functional consequences of mislocalizing PfSec14-2 and PfSec14-4, the PfSEC14-2-2xFKBP-mNG and PfSEC14-4-2xFKBP-mNG lines were transfected with a vector allowing the episomal expression of the FRB 3xNLS mCherry mislocalizer. Upon addition of rapalog, the FRB domain of the mislocalizer heterodimerizes with the FKBP domains of the tagged proteins, resulting in their nuclear translocation and preventing them from reaching their normal site of action, thereby causing a functional inactivation of the protein. Rapalog addition triggered efficient nuclear relocalization of both proteins (Fig. S6). Over a 120-hour time course, rapalog treatment did not measurably alter the PEMR relative to vehicle controls in either line. For PfSEC14-2 KS, the PEMR was 5.50 (SD 0.87) with rapalog versus 5.72 (SD 0.84) but this was not statistically significant (two-sided Mann–Whitney U test: p = 0.589; n = 3). For PfSEC14-4 KS, the PEMR was 7.28 (SD 2.61) with rapalog versus 6.63 (SD 2.26) with vehicle, a difference again not statistically different (two-sided Mann–Whitney U test: p = 0.310; n = 3) (Fig. 4D and Fig. S7). The absence of a significant growth defect upon induction of the KS in the PfSec14-2-2xFKBP-mNG/FRB-3xNLS-mCherry line is consistent with the TGD and the PiggyBac data (M. Zhang et al., 2018). However, the absence of a significant growth defect in the PfSec14-4-2xFKBP-mNG/FRB-3xNLS-mCherry line contrasts with our inability to generate a TGD line for this gene and with the PiggyBac data, which indicate that parasites with transposon insertions in the CDS or UTR regions of the gene could not be generated (M. Zhang et al. 2018). This discrepancy could be explained by an incomplete sequestration of the protein upon induction of the KS, with residual protein remaining at its normal site of action at levels sufficient to maintain its essential function.

Western blotting with an anti-HA antibody followed by densitometric analysis was performed in the PfSEC14-1^TetR^ line after induction of gene expression repression by removing anhydrotetracycline (aTC) from the culture medium, in order to quantify the extent of repression. An approximately 70% reduction in PfSec14-1-HA abundance was observed following aTC withdrawal (Fig. S5C). Despite this depletion, parasite growth over 120 hours was not detectably different between conditions: PEMR = 5.29 (SD 1.84) without aTC versus 5.10 (SD 2.08) with aTC (two-sided Mann–Whitney U test, p = 0.937; n = 3) (Fig. S5D and S5E). These data suggest that the extent of knockdown achieved here is insufficient to elicit a measurable phenotype.

Taken together, these data indicate that PfSec14-2 does not appear to be essential for normal asexual parasite growth. In this experimental context, PfSec14-4 also appears dispensable, which is at odds with our inability to generate a TGD line. This contradiction may reflect limitations inherent to the KS technique, which may fail to sequester 100% of the target protein and thus leave sufficient residual activity to support normal growth. The inability to recover TGD or KS lines for PfSEC14-3, PfSEC14-5, and PfSEC14-6 could indicate an important role in asexual proliferation but may also reflect locus-specific technical refractoriness to genome editing or intolerance to C-terminal tagging.

Comparing these observations with orthologs from other apicomplexans provides additional perspective. The pronounced growth defect observed for PfSEC14-1 TGD, together with our inability to generate a KS line, is consistent with the probable essentiality of the *P. berghei* ortholog (PBANKA_1125200) (Table 1) (Bushell et al. 2017) but does not clearly match the apparently dispensable phenotype reported for the *T. gondii* ortholog (TGME49_237000) (Sidik et al. 2016). For PfSEC14-3, the lack of recoverable mutants is broadly in line with the low fitness score caused by the disruption of its *T. gondii* ortholog (TGME49_315590) (Sidik et al. 2016). Regarding PfSEC14-5, our failure to recover TGD and KS lines contrasts with *P. berghei*, in which PBANKA_1205400 was dispensable for proliferation in mice but affected the sporozoite-to-blood-stage transition (Table 1) (Bushell et al. 2017), and with *T. gondii*, where loss of TGME49_233300 increased *in vitro* fitness (Ishizaki et al. 2022). Notably, truncation of the PfSEC14-5 C-terminal RhoGAP domain has been associated with increased merozoite invasiveness (K. Kumar et al. 2017); because our TGD design results in the disruption of both the Sec14 domain and the RhoGAP domain, these findings raise the possibility that the RhoGAP domain modulates invasion phenotypes, whereas the Sec14 domain may support a function that becomes limiting for viability when removed in *P. falciparum.* Finally, PfSEC14-6 was refractory to modification even though the corresponding *T. gondii* gene (TGME49_237000) carries a higher fitness score, suggestive of dispensability in that system (Sidik et al. 2016). Overall, these comparisons support the idea that some of the functions of Sec14-family proteins may be conserved across apicomplexans, while others may be context- or species-dependent. The expanded Sec14 domain-containing protein repertoire in *Toxoplasma* could provide functional redundancy that mitigates the impact of losing individual members, whereas *P. falciparum* encodes a more limited set.

### PfSec14-1, PfSec14-2, and PfSec14-4 exhibit distinct localization patterns

We next investigated the cellular localization of mNG tagged PfSec14-1, PfSec14-2, and PfSec14-4.

#### PfSec14-1 transitions from nuclear to perinuclear localization during schizogony

Despite the absence of detection of PfSec14-1-2xFKBP-mNG by Western blot, a clear mNG signal was observed by fluorescence microscopy, supporting the hypothesis that the lack of detection in the Western blot is due to the limited sensitivity of the anti mNG antibody used. PfSec14-1-2xFKBP-mNG was detected throughout the asexual blood-stage cycle (Fig. 5A). In rings and trophozoites, the fluorescence appeared broadly cytosolic and overlapped with the Hoechst-stained nucleus (Fig. 5Ai and Aii); strikingly, in early schizonts the signal was predominantly nuclear (Fig. 5Aiii). As schizogony progressed, the nuclear enrichment diminished and the signal redistributed into a distinct “I shaped” pattern that partially overlapped with the nucleus (Fig. 5Aiv). During late schizogony, PfSec14-1-2xFKBP-mNG relocalized to the cytoplasm, and foci in close proximity to the nucleus became visible within a subset of developing merozoites (Fig. 5Av).

**Fig. 5.**
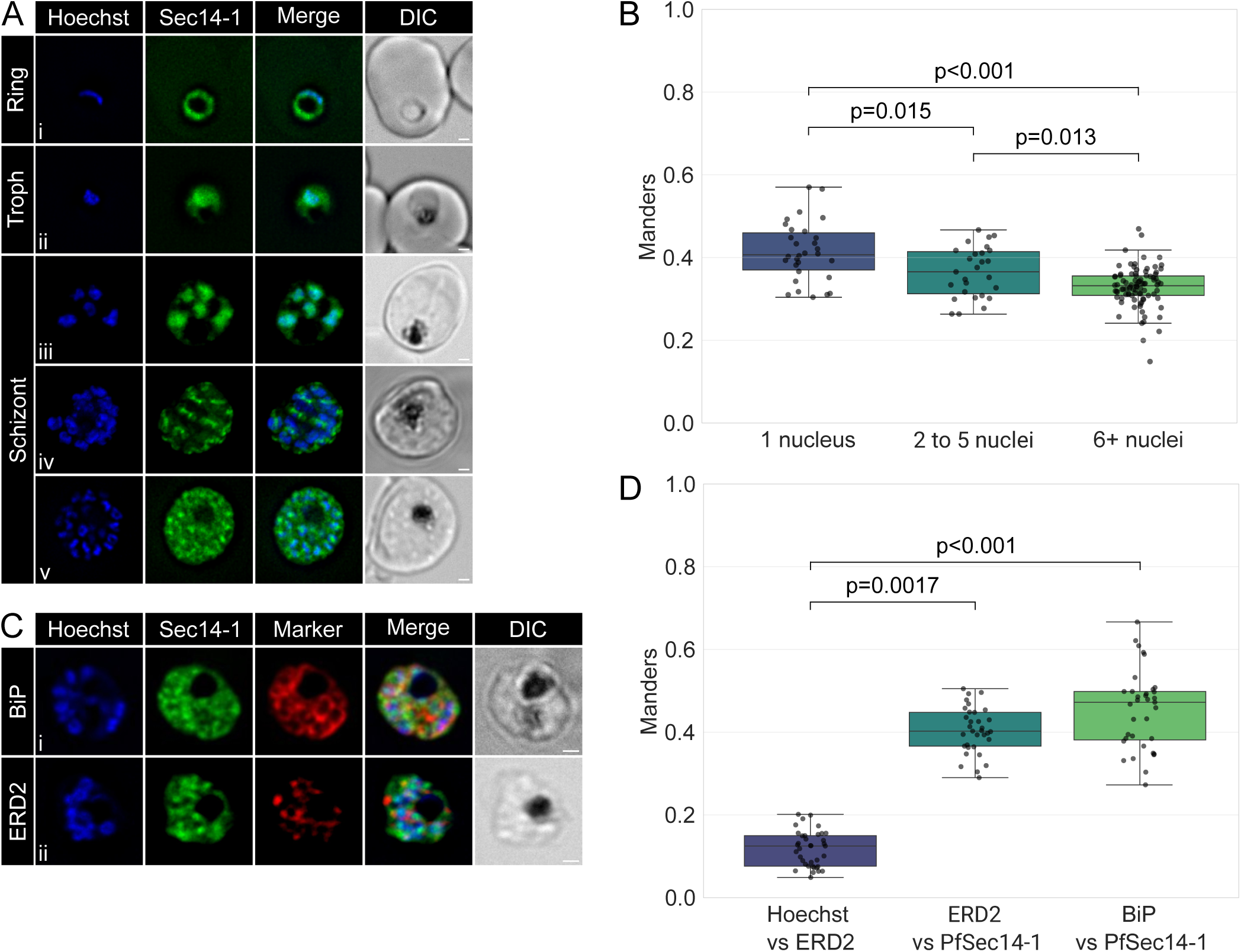
Localization of PfSec14-1. (A) Live-cell fluorescence microscopy showing the localization of PfSec14-1-2xFKBP-mNG throughout the asexual erythrocytic cycle. (B) Box plot showing the Mander’s overlap coefficient (M2) between Hoechst and PfSec14-1–2xFKBP–mNG signals, calculated according to the number of nuclei per parasite (cells with one nucleus, n = 30; cells with 2–5 nuclei, n = 27; cells with ≥6 nuclei, n = 90). (C) Live-cell fluorescence microscopy showing the expression of mNG-NLS^Sec14-1^. (D) Indirect immunofluorescence assay (IFA) on schizont-stage parasites to determine the overlap between PfSec14-1-2xFKBP-mNG and the endoplasmic reticulum marker BiP (Di) or the cis-Golgi marker ERD2 (Dii). (E) Box plots showing Mander’s overlap coefficients M2 (marker vs PfSec14-1-2xFKBP-mNG) between PfSec14-1-2xFKBP-mNG, and BiP (n=39) or ERD2 (n=33), Hoechst vs ERD2 Mander’s overlap coefficients was used as a negative control (n=37). All analyses were performed on images acquired from three biological replicates. Nuclei are stained with Hoechst (blue). Scale bar represent 1 µm. Only statistically significant differences between strains are indicated (Kruskal–Wallis test).

To quantify the stage dependent changes in nuclear localization suggested by these images, Manders’ overlap coefficients were calculated between the Hoechst-stained nuclei and the PfSec14-1-2xFKBP-mNG signal (Fig. 5B). Parasites were stratified into three developmental bins based on the number of nuclei per parasite (1 nucleus, 2–5 nuclei, and ≥6 nuclei), which serves as a proxy for developmental stage. A Kruskal–Wallis test indicated that Manders’ coefficients differed significantly across these groups (test statistic = 31.6348, p < 0.001). Pairwise two-sided Mann–Whitney U tests showed that colocalization was highest in the earliest stage: the 1-nucleus group (Manders’ coefficients ± SD = 0.4143 ± 0.0719, n = 30) exhibited significantly greater nuclear overlap than the 2–5 nuclei group (0.3644 ± 0.0608, n = 27; p = 0.015) and the ≥6 nuclei group (0.3286 ± 0.0479, n = 90; p < 0.001). Colocalization also differed significantly between the 2–5 nuclei and ≥6 nuclei groups (p = 0.013). Collectively, these results support a progressive reduction in PfSec14-1-2xFKBP-mNG nuclear localization as parasites advance from early to later developmental stages, consistent with the shift from nuclear enriched to perinuclear and cytoplasmic focused signal observed by fluorescence microscopy.

To characterize the perinuclear localization observed in schizonts, immunofluorescence assays (IFA) were performed using antibodies against the endoplasmic reticulum (ER) marker BiP (Fig. 5Ci) and the Golgi marker ERD2 (Fig. 5Cii). Colocalization analysis revealed partial overlap between PfSec14-1-2xFKBP-mNG and both the Golgi (Manders’ coefficient ± SD = 0.40 ± 0.06) and the ER (0.43 ± 0.11 SD) (Fig. 5D). As a negative control, Manders’ coefficients were calculated between the nuclear stain (Hoechst) and the Golgi marker (ERD2), yielding a low value (0.12 ± 0.04). Pairwise comparisons using the Kruskal–Wallis test indicated significantly higher colocalization of PfSec14-1-2xFKBP-mNG with the Golgi (Hoechst/ERD2 vs. ERD2/PfSec14-1, p < 0.001) and with the ER (Hoechst/ERD2 vs. BiP/PfSec14-1, p < 0.001).

PfSec14-1 displays a dynamic, developmentally regulated localization pattern across the *P. falciparum* erythrocytic cycle, shifting from prominent nuclear accumulation in trophozoites and early schizonts to a perinuclear distribution during merozoite maturation. This nuclear localization is reminiscent of the yeast Sec14-family homolog Sfh1p, which contains a predicted NLS and has been demonstrated to associate predominantly with the nucleus (Schnabl et al. 2003). Although the physiological function of Sfh1p’s nuclear localization remains incompletely understood, nuclear PIPs are now recognized as critical regulators of gene expression, chromatin remodeling, and nuclear signaling in diverse eukaryotic systems (Vidalle et al. 2023). The pronounced nuclear accumulation during trophozoite and early schizont stages, periods marked by intensive transcriptional activity and DNA replication, raises the possibility that PfSec14-1 supplies PI to support nuclear PIPs synthesis for chromatin organization (Vidalle et al. 2023) or stage-specific transcriptional regulation.

The localization of PfSec14-1-2xFKBP-mNG to the ER/Golgi in late schizonts also suggests a potential functional connection with the PI4-kinase PfPI4KIIIβ. *P. falciparum* encodes three putative PI4-kinases, and inhibition of the PfPI4KIIIβ homologue by imidazopyrazines and quinoxalines causes a striking block in cytokinesis, characterized by defects in membrane biogenesis and ingression around developing merozoites (McNamara et al. 2013). In line with this, PI4P has been mapped to the Golgi and plasma membrane in blood-stage parasites (Ebrahimzadeh, Mukherjee, and Richard 2018), and PfPI4KIIIβ itself shifts from a diffuse cytosolic distribution in trophozoites to discrete foci in schizonts that likely correspond to the Golgi apparatus (McNamara et al. 2013). Moreover, our data show that PfSec14-1 can partially rescue the yeast *sec14-1^ts^* growth defect, suggesting PI/PC exchange activity. It is therefore tempting to speculate that late-stage Golgi-associated PfSec14-1 could act upstream of PfPI4KIIIβ by supplying PI to Golgi membranes, thereby supporting local PI4P production required for vesicular trafficking and membrane ingression during merozoite formation.

#### Stage specific recruitment of PfSec14-2 to the plasma membrane, inner membrane complex and basal complex

We next examined the localization of PfSec14-2-2xFKBP-mNG. Fluorescence was weak and diffusely cytosolic during ring stages (Fig. 6Ai). From trophozoites (Fig. 6Aii) through early schizonts (Fig. 6Aiii), the signal remained faint but began accumulating in discrete foci. In mid-schizonts (Fig. 6Aiv), it adopted a ring-like distribution, which transitioned in late schizonts (Fig. 6Av) to a more linear structure oriented parallel to the RBC membrane within each developing merozoite, consistently positioned posterior to the nucleus when viewed relative to the host membrane boundary. In parasites arrested prior to egress with ML10 (Fig. 6Avi), a highly specific inhibitor of parasite cGMP-dependent protein kinase (PKG) that reversibly blocks schizont rupture (Ressurreição et al. 2020), PfSec14-2-2xFKBP-mNG signal intensity diminished markedly, with only faint foci detectable. This ring-like pattern in mid/late schizogony that evolves into foci resembles the localization dynamics previously reported for the basal complex marker PfPPP8 (Morano, Rudlaff, and Dvorin 2023).

**Fig. 6.**
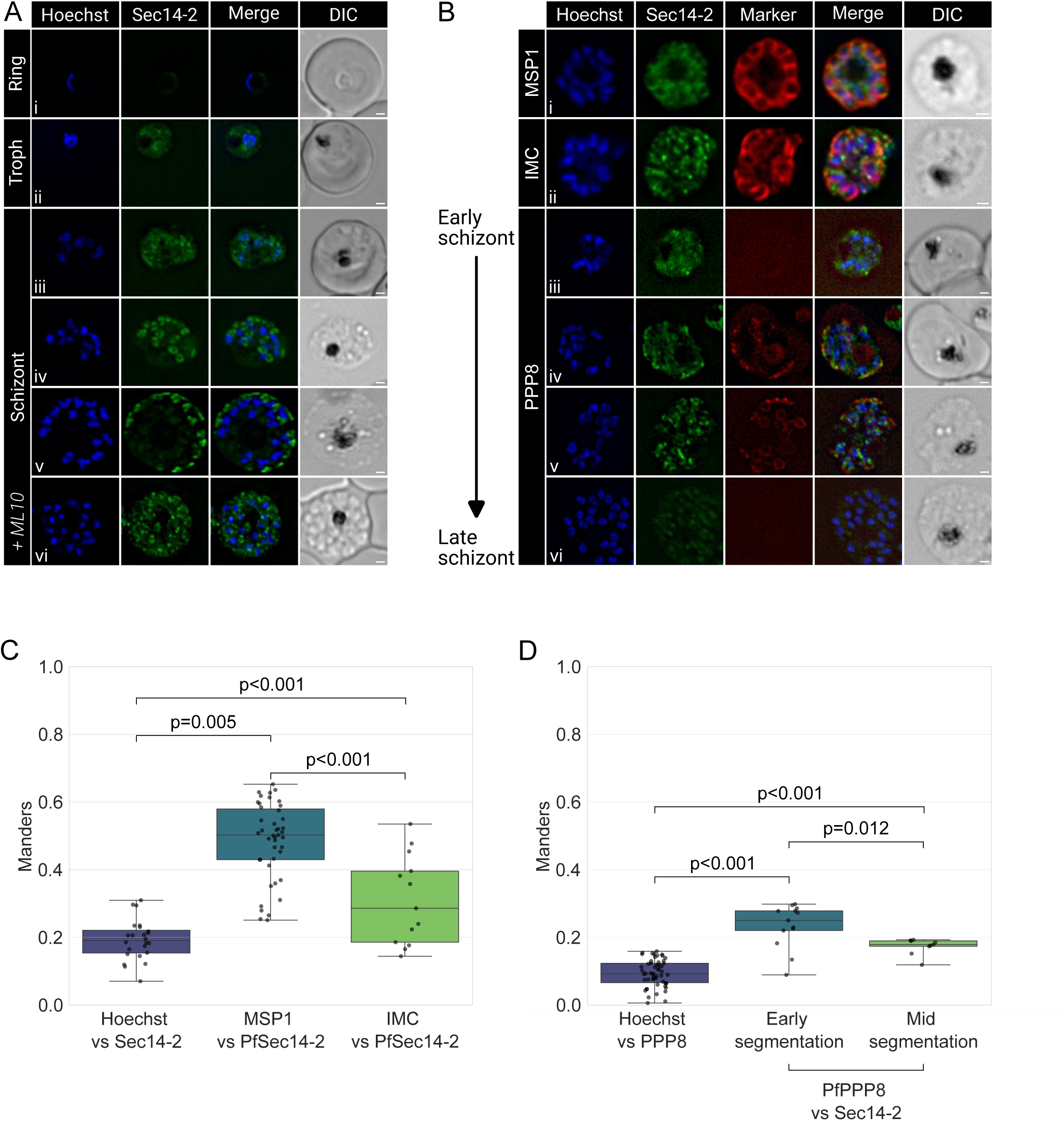
Localization of PfSec14-2. (A) Live-cell fluorescence microscopy showing the localization of PfSec14-2-2xFKBP-mNG across the asexual erythrocytic stages. (B) IFA on PfSec14-2-2xFKBP-mNG schizont-stage parasites using anti-MSP1 (Bi) and anti-IMC antibodies (Bii). Live-cell fluorescence microscopy showing expression of PfSec14-2-2xFKBP-mNG/PfPPP8-mCh during schizogony (Biii - Bvi). (C) Box plots showing Mander’s overlap coefficients M2 (marker vs PfSec14-2-2xFKBP-mNG) between PfSec14-2-2xFKBP-mNG, the IMC (n=13) and MSP1 (n=43), Hoechst vs ERD2 Mander’s overlap coefficients was used as a negative control (n=24). (D) Box plots showing Mander’s overlap coefficients M2 (marker vs PfSec14-2-2xFKBP-mNG) between PfSec14-2-2xFKBP-mNG and PfPPP8 during segmentation (early segmentation cells, n=13; mid segmentation cells, n=9), Hoechst vs PfPPP8 Mander’s overlap coefficients M2 was used as a negative control (n=54). All analyses were performed on images acquired from three biological replicates. Nuclei are stained with Hoechst (blue). Scale bars represent 1 µm. Only statistically significant differences between strains are indicated (Kruskal–Wallis test).

IFA with an anti-MSP1 antibody, a parasite plasma membrane (PPM) marker (Fig. 6Bi), revealed partial colocalization of PfSec14-2-2xFKBP-mNG with the PPM (Manders’ coefficient ± SD = 0.48 ± 0.11), whereas colocalization with the inner membrane complex (IMC) marked using an anti-IMC antibody (Fig. 6Bii) was weaker (0.31 ± 0.11 SD) (Fig. 6C). Hoechst nuclear staining served as a negative control, yielding minimal colocalization (0.19 ± 0.06). Kruskal–Wallis analysis confirmed statistically significant enrichment of PfSec14-2-2xFKBP-mNG signal at the PPM (p = 0.005) and IMC (p < 0.001) relative to the nucleus (Fig. 6C).

To probe a putative basal complex association, we generated a dual-labeled line by transfecting PfSEC14-2-2xFKBP-mNG parasites with a vector enabling endogenous mCherry-tagging of PfPPP8 (PfSEC14-2-2xFKBP-mNG/PfPPP8-mCherry). PfPPP8 is a basal complex–associated pseudophosphatase that is essential for maintaining its integrity during schizogony. As a *bona fide* basal complex protein, loss of PfPPP8 results in fragmentation of the basal complex ring. PfPPP8 also displays a distinct temporal localization pattern: it is recruited to the basal complex during its expansion phase but dissociates before basal complex contraction and completion of cytokinesis (Morano, Rudlaff, and Dvorin 2023). In the PfSEC14-2-2xFKBP-mNG/PfPPP8-mCherry line, the two signals partially overlapped during early segmentation (Fig. 6Biv) (Manders’ coefficient ± SD = 0.23 ± 0.06). In mid-segmentation (Fig. 6Bv), the overlap was less pronounced (0.17 ± 0.02). By late segmentation, as parasites approached egress, the PfPPP8 signal was no longer detectable, while the PfSEC14-2-2xFKBP-mNG signal became faint with residual foci (Fig. 6Bvi). Hoechst nuclear staining, used as a negative control, showed minimal colocalization with PfPPP8 (0.09 ± 0.04). Kruskal–Wallis analysis confirmed significant colocalization of PfPPP8-mCherry with PfSEC14-2-2xFKBP-mNG during early (p < 0.001) and mid-segmentation (p < 0.001) relative to the nucleus (Fig. 6D). Furthermore, the degree of colocalization was significantly reduced in mid-segmentation compared with early segmentation (p = 0.012).

It was previously shown that PfSec14-2 exhibits cholesterol, PI(4,5)P, and PC transfer activities in liposome-based in vitro assays using radiolabeling, fluorescence resonance energy transfer, fluorescence cross-correlation spectroscopy, and fluorescence dequenching assays with recombinant protein (Šťastný et al. 2025). In *P. falciparum*, PI(4,5)P is present at the PPM (Ebrahimzadeh, Mukherjee, and Richard 2018) and although cholesterol has been detected at the IMC in *T. gondii* (Coppens and Joiner 2003; Johnson et al. 2007), its presence at the *P. falciparum* IMC remains unknown. The IMC protein PfIMC1l binds both cholesterol and PI(4,5)P *in vitro* (V. Kumar et al. 2019), suggesting that these lipids might be present at the IMC. The spatiotemporal dynamics of PfSec14-2 at the PPM, IMC, and basal complex (mirroring the expression pattern of the basal complex marker PfPPP8) together with its lipid-transfer activity, are consistent with a potential role in supporting local lipid homeostasis during IMC assembly. Given the tight apposition of the PPM, IMC, and basal complex in late schizonts, colocalization analysis warrants cautious interpretation. Nonetheless, PfSec14-2’s *in vitro* lipid-transfer activity and spatiotemporal localization pattern are consistent with a role in local lipid exchange between the PPM and adjacent IMC, potentially modulating membrane composition and effector recruitment (e.g., PfIMC1l) during IMC biogenesis. The absence of growth defects in PfSEC14-2 KS and TGD lines during the intraerythrocytic cycle indicates dispensability for asexual blood-stage proliferation, potentially due to redundancy with other lipid-transfer or trafficking pathways. Notably, PfIMC1l has been proposed to contribute to sporozoite motility through potential interactions linking the IMC with the glideosome machinery (V. Kumar et al. 2019), suggesting PfSec14-2 may play a more prominent role in sporozoites.

#### PfSec14-4 localizes to the ER/Golgi in schizonts

PfSEC14-4-2xFKBP-mNG displays weak, predominantly cytoplasmic fluorescence in ring stages (Fig. 7Ai). In trophozoite the signal is almost absent (Fig. 7Aii), similarly to what observed by Western blot. As schizogony progresses, the signal intensifies and adopts a characteristic perinuclear ring-like pattern reminiscent of ER labeling (Fig. 7Aiii to 7Av). IFA performed on schizonts using ER (BiP) (Fig. 7Bi) and Golgi (ERD2) (Fig. 7Bii) markers revealed partial colocalization with both compartments (Fig. 7C). The Manders’ coefficient (± SD) between BiP and PfSec14-4-2xFKBP-mNG was 0.44 ± 0.10 whilst it was 0.43 ± 0.11 between ERD2 and PfSec14-4-2xFKBP-mNG. The Manders’ coefficient between Hoechst and ERD2 was used as a negative control (0.14 ± 0.10). A Kruskal–Wallis analysis confirmed these values significantly exceed the negative control (PfSec14-2-2xFKBP-mNG/BiP: p < 0.001; PfSec14-2-2xFKBP-mNG/ERD2: p < 0.001).

**Fig. 7.**
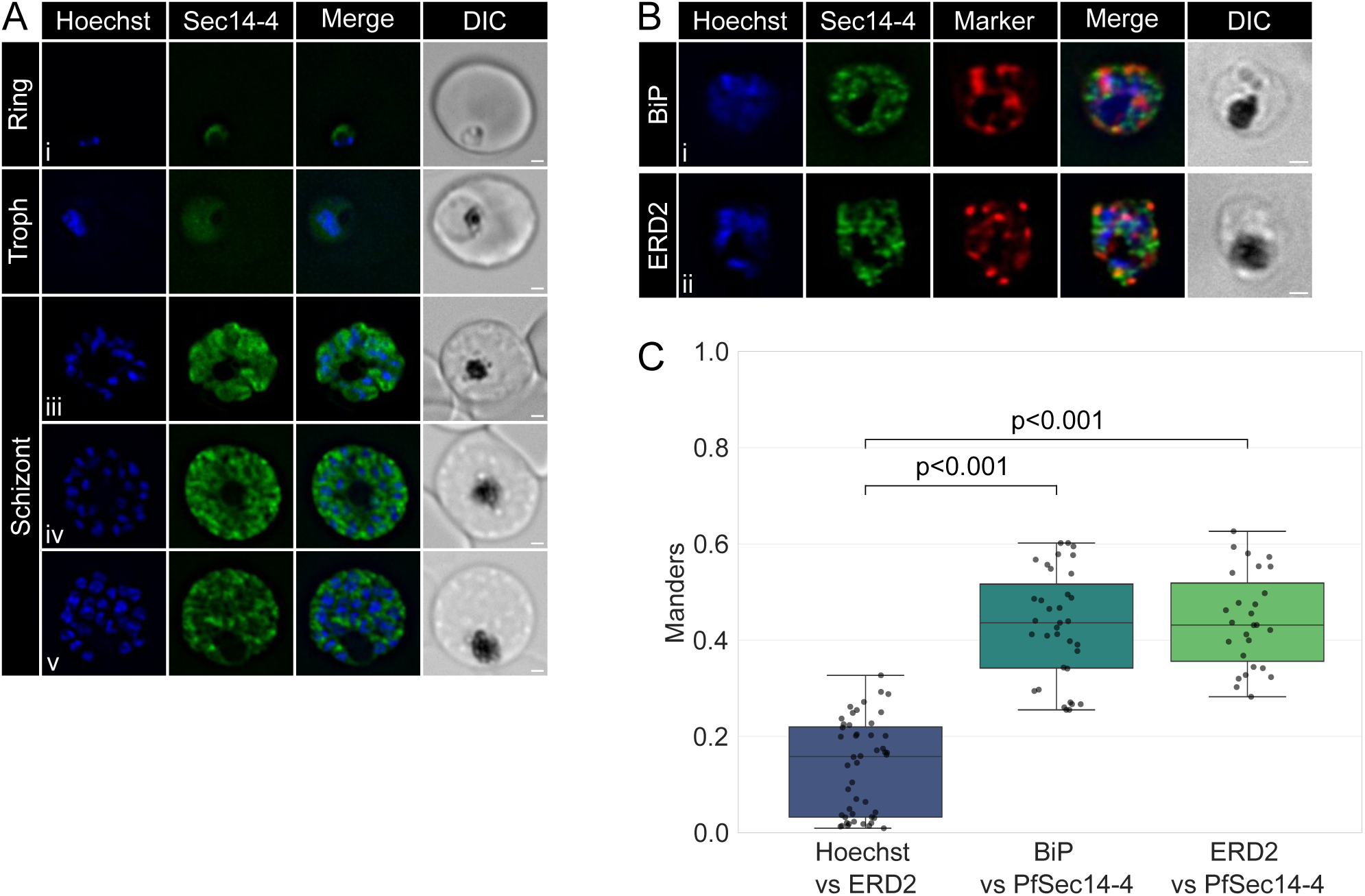
Localization of PfSec14-4. (A) Live-cell fluorescence microscopy showing the localization of PfSec14-4-2xFKBP-mNG across the asexual erythrocytic stages. (B) IFA on schizont-stage parasites to determine the overlap between PfSec14-4-2xFKBP-mNG and the endoplasmic reticulum marker BiP (Bi) or the cis-Golgi marker ERD2 (Bii). (C) Box plots showing Mander’s overlap coefficients M2 (marker vs PfSec14-4-2xFKBP-mNG) between PfSec14-4-2xFKBP-mNG, and the BiP (n=35) or ERD2 (n=27), Hoechst vs ERD2 Mander’s overlap coefficients was used as a negative control (n=48). All analyses were performed on images acquired from three biological replicates. Nuclei are stained with Hoechst (blue). Scale bar represent 1 µm. Only statistically significant differences between strains are indicated (Kruskal–Wallis test).

Overall, PfSec14-4-2xFKBP-mNG localizes to a perinuclear ring-like structure in schizonts, primarily colocalizing with the ER marker BiP and showing discrete foci overlapping the Golgi marker ERD2 similar to PfSec14-1-2xFKBP-mNG in late schizonts. Given PfSec14-4’s partial complementation of yeast Sec14p (suggesting possible PI/PC transfer activity, akin to PfSec14-1), it might facilitate lipid exchange between ER and Golgi to support PIP synthesis. Both proteins’ shared localization and apparent transfer activities could imply functional redundancy during schizogony, potentially allowing mutual compensation; this might explain why PfSec14-4 KS has no growth impact, while PfSec14-1 disruption causes a strong defect, possibly due to PfSec14-1’s unique early nuclear localization that PfSec14-4 lacks.

It would have been interesting to induce the KS of PfSec14-1 specifically in mid-schizonts to determine whether its absence at this stage affects parasite development. If no effect were observed, it would suggest that the nuclear presence of PfSec14-1 is uniquely important and cannot be compensated for by other Sec14 proteins that do not localize to the nucleus. Unfortunately, our inability to generate a pure PfSEC14-1-2xFKBP-mNG line prevented us from performing this experiment.

## Discussion

Targeted disruption of PfSEC14-1 demonstrates that PfSec14-1 is likely important, as evidenced by the severe growth defect in the TGD line. In contrast, PfSEC14-2 is dispensable, with KS and TGD lines showing normal proliferation. The successful generation of a KS line for PfSEC14 4 and the absence of growth defects upon induction of PfSec14 4 mislocalization contradict the apparent essentiality suggested by our inability to generate a TGD line. However, this lack of phenotype could result from incomplete mislocalization, leaving sufficient protein at its site of action to maintain normal function. Finally, PfSEC14-3, -5, and -6 appear refractory to modification, suggesting essentiality.

mNG tagging enabled localization of PfSec14-1, -2, and -4, revealing distinct patterns: PfSec14-1 shows dynamic nuclear to ER/Golgi localization, PfSec14-2 seems to potentially associate with PPM/IMC/basal complex, and PfSec14-4 localizes to the ER/Golgi. These three proteins therefore look to be localizing to membranes of distinct organelles. Given that canonical Sec14 proteins transfer lipids between membranes, these localization patterns are consistent with a potential LTP function for these proteins.

Yeast complementation suggests PI/PC-transfer activity for PfSEC14-1, -4, and -6 (partial *sec14^ts^*rescue), supporting their function as PITPs in *P. falciparum.* Nonetheless, the fact that these three proteins might share a similar PI/PC transfer activity suggests potential functional redundancy, which could explain, for example, why the PfSec14 1 TGD line exhibits a slow growth phenotype but not lethality, potentially due to partial compensation by PfSec14 4, which shares a common ER/Golgi localization in late stage parasites.

Numerous questions about Sec14 function remain unresolved, and paralog redundancy complicates single-gene analyses (Huang, Ghosh, and Bankaitis 2016). Future studies in *P. falciparum* should pursue multi-gene conditional KD or KO, though this is currently limited by the few available selectable markers (Mogollon et al. 2016). To better define substrate specificities, *in vitro* lipid-binding assays should be extended to the remaining PfSec14 proteins, building on prior characterization of PfSEC14-2 and PfSEC14-6 (Šťastný et al. 2025). Domain-specific truncation studies would further dissect multifunctional Sec14 proteins. For multidomain members like PfSEC14-3 (Macro) and PfSEC14-5 (RhoGAP), targeted deletions could isolate Sec14 versus second domain contributions to essentiality. Similarly, truncating the large uncharacterized C-terminal domain of PfSEC14-1 could test its role in nuclear targeting/retention, potentially unmasking separable functions currently obscured by full-length disruption phenotypes. A stage-specific PfSEC14-1 knockdown line (attempted using both KS and TetR systems but unsuccessful in yielding a usable line) could help dissect the respective nuclear and ER/Golgi functions of this protein. Additionally, the KS lines should be tested in other life-cycle stages (hepatic, sexual, and mosquito) to determine stage-specific essentiality and potential transmission-blocking activity.

Ultimately, Sec14 PITPs represent druggable targets with species-selective inhibitors proven in fungi (Chen et al. 2023). The putative essential roles of some Sec14 domain-containing proteins in *P. falciparum* underscore therapeutic potential against malaria.

## Experimental Procedures

### Discrimination of Sec14 and BCH domain families in predicted *P. falciparum* Sec14 proteins

Protein domain architectures and domain positions were further analyzed using the InterPro database (https://www.ebi.ac.uk/interpro/) (Blum et al. 2025). To distinguish if any proteins contained BCH domains, a subclass of the Sec14 superfamily (Gupta et al. 2012), hidden Markov model (HMM) profiles were generated for BCH and Sec14 domains using the GENOME.jp motif tool(Eddy 1998). For this, we aligned sequences of 175 putative BCH-domain proteins and 69 Sec14-domain proteins (Gupta et al. 2012) using MUSCLE v5.2 (Edgar 2022). The resulting profiles were then applied to the *P. falciparum* 3D7 RefSeq protein database to identify Sec14 and BCH domains.

### Proteins structure prediction and structural alignment

The 3D structures of the Sec14 domains were predicted using the AlphaFold server (https://alphafoldserver.com), implementing the AlphaFold 3 algorithm (Abramson et al. 2024). For each protein, only the Sec14 domain sequence was input (amino acids 49–281 for PfSec14-1, 770–908 for PfSec14-3, 215–542 for PfSec14-4, and 29–159 for PfSec14-5). Full-length sequences were used for PfSec14-2 and PfSec14-6. Structural comparisons were performed using US-align (version 20240510) (C. Zhang et al. 2022) with default parameters for sequence-independent alignment. TM-score statistics quantified structural similarity. Predicted Sec14 domain proteins were compared with the *S. cerevisiae* Sec14p crystal structure (PDB: 7ZG9) (Chen et al. 2023), while predicted BCH-domain proteins were compared to the *S. pombe* BCH domain structure (PDB: 7E0W) (Chichili et al. 2021).

### Yeast strains and culture conditions

*S. cerevisiae* strains used for functional complementation were as follows: the *sec14^ts^* strain (**a** *leu2*, *trp1*, *lys2*, *ura3*, *his3*, *sec14-1^ts^*) is a spore from a genetic cross of CTY1-1A strain (kindly provided by V. Bankaitis, Texas A&M University, USA) (Fang et al. 1996) and a yeast strain of w303 origin (_α_ *his3*, *leu2*, *trp1*, *ura3*). Azole sensitive *pdr16*::*hisG* strain (**a** *ura3-52*, *leu2*Δ*1*, *his3*Δ*200*, *trp1*Δ*63*, *GAL2*,*pdr16*_∷_*hisG*) in the FY1679-28c genetic background, originally from A. Goffeau laboratory (Catholic University Louvain, Belgium) (van den Hazel et al. 1999) was kindly provided by G. Daum (Technical University, Graz, Austria). *S. cerevisiae psd1*_∷_*TRP1 pdr17*_∷_*HIS3* strain WWY62 (**a** *lys2*, *ura3*, *his3*, *leu2*, *trp1*, *met*, *psd1*_∷_*TRP1*, *pdr17*_∷_*HIS3*) used to study complementation of the absence of Pdr17p is a gift by D. Voelker (National Jewish Health, Denver, CO, USA) (Trotter et al. 1998). Yeast strain to explore complementation of the absence of all yeast Osh proteins JRY6326 (_α_ *leu2-3,112*, *ura3-52*, *his3*Δ*200*, *trp1-*Δ*901*, *suc2*Δ*9*, *osh1*Δ::*kanMX4*, *osh2*Δ::*kanMX4*, *osh3*Δ::*LYS2*, *osh4*Δ::*HIS3*, *osh5*Δ::*LEU2*, *osh6*Δ::*LEU2*, *osh7*Δ::*HIS3*, *p^MET^- OSH2-TRP1*) was kindly provided by C. Beh (Simon Fraser University, Canada) (Beh et al. 2001).

Yeast strains were grown on yeast extract/peptone/dextrose (YEPD; 2 % glucose) media unless otherwise stated. Media for strains required ethanolamine were supplemented with 2 mM ethanolamine after sterilization. The *osh1*Δ *- 7*Δ *p^MET3^-OSH2* strain was grown on standard synthetic minimal medium (0.67 % YNB without amino acids, 2 % glucose) without methionine (YNB-MET). Yeast strains containing episomal plasmids were maintained and pre-grown on standard synthetic minimal media (0.67 % YNB without amino acids, 2 % glucose) supplemented with essential amino acids and bases as required for plasmid maintenance. Solid media were supplemented with 2 % agar.

### Yeast phenotypic testing

To explore complementation of *S. cerevisiae* phenotypes, the coding sequences of Sec14 domain-containing genes of *P. falciparum* were codon-optimized for yeast expression and commercially synthesized by genscript. The synthetic sequences were digested with *HindIII* and *BamHI* and cloned into the p416-GPD plasmid (Mumberg, Müller, and Funk 1995) under the control of the strong constitutive glyceraldehyde-3-phosphate dehydrogenase gene (GPD) promoter (Blazeck et al. 2012), and C-terminally tagged with a Myc epitope. Due to gene size and sequence complexity, only the predicted Sec14 domains sequence of *PfSec14-1* and *PfSec14-4* were synthesized while the full-length (FL) sequence was used for the 4 other genes. Complementation of yeast phenotypes by *P. falciparum* Sec14 domain-containing proteins were performed by spot assays. Yeast cultures were pre-grown overnight in media appropriate for the maintenance of plasmids. Yeast cultures were spotted as 10-fold dilutions. The growth was scored after 3 days of incubation at indicated temperatures.

### Parasite culture

*P. falciparum* 3D7 and NF54 asexual blood-stage parasites (from David Walliker, Edinburgh University) were cultured under standard conditions in RPMI-HEPES medium with 4% hematocrit (human O+ erythrocytes) and 0.5% (w/v) Albumax™ (Invitrogen) at 37°C in a controlled gas mixture (5% O, 5% CO, 90% N) (Trager and Jensen 1976).

### Vector construction and transfection

Primers used for all constructs are listed in S1 Table.

#### Targeted Gene Disruption (TGD) Constructs

To disrupt the genes of interest (GOI), we used the selection-linked integration (SLI) method (Birnbaum et al. 2017). Approximately 500 bp from the N-terminus of each GOI was PCR-amplified using primers harboring 5’ NotI and 3’ MluI restriction sites. The amplicons were cloned in-frame with GFP into pSLI-TGD vector digested with NotI and MluI. To prevent expression of the disrupted locus, the start codon (ATG) was replaced with a stop codon (TAA) in the 5’ primer sequence (Birnbaum et al. 2017).

#### Knock-Sideways (KS) Constructs

To replace GFP with mNeonGreen in the pSLI-2×FKBP-GFP vector(Birnbaum et al. 2017), the mNeonGreen (mNG) sequence was amplified using primers containing 5′ BsiWI and 3′ SalI restriction sites and cloned in-frame into the pSLI-2×FKBP-GFP vector digested with BsiWI and SalI. The neomycin resistance cassette was then replaced with yDHODH by amplifying the yDHODH sequence with primers containing 5′ T2A-SalI and 3′ XhoI sites and cloning it in-frame into the pSLI-2×FKBP-mNG vector digested with SalI and XhoI.

GOIs were endogenously tagged at the C terminus with 2×FKBP-mNG using SLI (Birnbaum et al. 2017). Approximately 500 bp of the GOIs’ C terminal region were PCR-amplified from *P. falciparum* genomic DNA with primers containing 5′ NotI and 3′ AvrII restriction sites and cloned in-frame into the pSLI-2×FKBP-mNG vector digested with NotI and AvrII.

#### TetR DOZI Conditional Knockdown vector

The pKD^PfSec14-1 repair template was assembled via NEBuilder® HiFi DNA Assembly, combining 5’HR and 3’HR homology arms amplified from *P. falciparum* 3D7 genomic DNA into AscI- and AatII- digested pKD vector (Rajaram, Liu, and Prigge 2020). The pH-gC^PfSec14-1 vector expressing Cas9 and gRNA was generated by cloning guide RNA sequences, selected using the online tool CRISPOR (Concordet and Haeussler 2018), into BsaI-digested pH-gC plasmid (Filarsky et al. 2018).

### Parasite transfection

Parasite transfections were performed following established protocols with slight modifications (Birnbaum et al. 2017). Briefly, *P. falciparum* 3D7 or NF54 parasites were transfected with 100 µg of pSLI-TGD (3D7) or pSLI-2×FKBP (NF54) constructs, or with 75 µg each of pKD^PfSec14-1^ and pH-gC^PfSec14-1^ vectors (3D7). Subsequent transfections were carried out in pSLI-Sec14-2-2×FKBP-mNG and pSLI-Sec14-4-2×FKBP-mNG lines using the mislocalizer plasmid p3×NLS-FRB-mCherry (Birnbaum et al. 2017) to enable knock-sideways induction. To generate the PfSec14-2-2×FKBP-mNG / PfPPP8-mCherry line, PfSec14-2-2×FKBP-mNG parasites were transfected with 75 µg of the PfPPP8-mCherry template vector and 75 µg of the pRR221 Cas9/gRNA expression vector (Morano, Rudlaff, and Dvorin 2023).

Transfectants were selected with 5 nM WR99210 for initial positive selection, followed by either 400 µg mL^-1^ neomycin (for pSLI-TGD) or 1.5 µM DSM1 (for pSLI-2×FKBP). WR99210 was reintroduced after parasite reemergence (∼10 days). For the PfSec14-1^TetR^ strain, 0.5 µM anhydrotetracycline (aTC) was maintained for 48 h, after which cultures were treated with 200 µg mL^-1^ blasticidin S-HCl and 1.5 µM DSM1 in the presence of aTC for 7 days. After selection, drug pressure was removed while maintaining aTC. Blasticidin S-HCl (200 µg mL^-1^) was also used to select for lines carrying the p3×NLS-FRB-mCherry or pHSP86-NLS^PfSec14-1^-mNG episomal vectors, as well as for PfPPP8-mCherry integration. Single clones were isolated by plaque assay(Thomas et al. 2016), and correct genomic integration was verified by diagnostic PCR (Table S1).

### Western blotting

To follow expression of *P. falciparum* Sec14-like proteins in *S. cerevisiae*, *sec14-1^ts^* yeast strain with respective p416-GPD-*Pf* Sec14-X-myc plasmids was pre-grown in YNB-URA medium at 26 °C. The next day the cultures were diluted and grown for additional 6 hours in YNB-URA at 26 °C to reach the exponential phase of growth (between 1 – 2.5 x 10^7^ cells/ml). Cells were disrupted by glass beads in the presence of protease inhibitors, cell homogenates were centrifuged, equivalent of 50 μg of protein was precipitated by TCA, mixed with loading buffer, and used for SDS-PAGE and subsequent western blotting. Primary antibody was THE™ c-Myc Tag Antibody, mAb, Mouse (GenScript, A007004-100) used in 1:1000 dilution, secondary antibody was Anti mouse IgG HRP conjugate (Sigma, 12349), used in 1:40000 dilution.

Parasite cultures were synchronized by two sorbitol treatments performed 18 hours apart, resulting in tightly synchronized parasites (18–24 h post invasion). Samples were harvested at the ring (12–18 h), trophozoite (26–32 h), and schizont (40–46 h) stages. Parasite pellets were lysed with saponin, solubilized in SDS sample buffer, separated by SDS-PAGE, and transferred onto PVDF membranes. Detection was carried out with HRP-conjugated secondary antibodies and enhanced chemiluminescence (ECL; Azure Biosystems). Equivalent parasite numbers were loaded for comparative expression analysis.

Dilutions for primary antibodies were as follows: 1:2500 for rabbit anti-mNeonGreen (proteintech, 29523-1-AP), 1:2000 for mouse anti-HA (Cederlane, CLH104AP) and 1:5000 for rabbit anti-PfHSP70 (StressMarq, SPC-186C).

### Growth assays

Synchronization was achieved by Percoll purification of schizonts, which were then cultured with uninfected red blood cells for 4 hours to allow merozoite invasion. Cultures were subsequently treated with sorbitol to retain only 0–4 h ring stage parasites. Parasites were then plated at 0.2% parasitemia. KS strain cultures were maintained either in the presence of 100 nM A/C heterodimerizer (Takara, 635056) or with ethanol vehicle as a control. The PfSEC14-1^TetR^ strain was cultured with 0.5 µM anhydrotetracycline (Cayman Chemical, 10009542) or with DMSO vehicle. Samples were collected at 24, 72, and 120 h intervals, stained with SYBR Gold (Invitrogen), and fixed with 1% paraformaldehyde. Parasitemia was quantified by flow cytometry using a BD FACSCanto II system operated with FACSDiva software, and data were analyzed with the flowkit Python package (White et al. 2021). Uninfected red blood cells were used to establish FITC thresholds. Statistical significance was evaluated using the Mann–Whitney U test implemented in the SciPy library (Virtanen et al. 2020). The parasite erythrocyte multiplication rate (PEMR) was calculated by dividing the parasitemia of the ring-stage culture by that of the culture 48 h earlier, corresponding to one complete replication cycle.

### Microscopy

Fluorescence microscopy acquisition was performed as previously described (Hallée and Richard 2015) using a GE Applied Precision Deltavision Elite microscope with a 100× 1.4 NA objective and with a sCMOS camera and deconvolved with the SoftWorx software.

Immunofluorescence assays (IFA) were performed following a previously described protocol (Mehnert, Simon, and Guizetti 2019), with minor modifications. Briefly, microscope slides wells were delimited using a DAKO hydrophobic barrier pen. Each well was coated with 100 µL Concanavalin A (Sigma–Aldrich; 5 mg mL^-1^ in water) and incubated for 20 min at 37°C, followed by two rinses with pre-warmed complete culture medium. Infected red blood cells (iRBCs) were pelleted by centrifugation, resuspended in pre-warmed complete medium at 0.03% hematocrit, and 100 µL of culture was added per well. Cells were allowed to settle for 10 min at 37°C, and unbound erythrocytes were removed by three gentle washes with pre-warmed medium. Cells were quickly rinsed with PBS and fixed with pre-warmed 4% paraformaldehyde (PFA) at 37°C for 20 min. Subsequent steps were carried out at room temperature. Samples were permeabilized with 0.1% Triton X-100 in PBS for 15 min and washed three times with PBS. To quench free aldehyde groups, a 0.1% (w/v) sodium borohydride (NaBH) solution was freshly prepared (0.01 g NaBH in 10 mL PBS) and 100 µL per well was added for a 10 min incubation. Slides were washed twice with PBS and then blocked with 3% BSA in PBS for 30 min. Primary antibodies were diluted in blocking solution and added at 100 µL per well for a 1 h incubation at room temperature. After three washes with 0.5% Tween 20 in PBS, secondary antibodies and Hoechst 33342 were diluted in blocking solution and incubated for 30 min. Samples were washed three times with 0.5% Tween 20 in PBS. Following staining, slides were mounted with antifade mounting medium and overlaid with coverslips.

Dilutions for primary antibodies were as follows: 1:2000 for rabbit anti-ERD2 (BEI Resources, MRA-72), 1:500 for rabbbit anti-BiP (Absalon, Robbins, and Dvorin 2016), 1:1000 for rabbit anti-IMC (Cepeda Diaz et al. 2023) and 1:2000 for rabbit anti-MSP1-19 (Blackman et al. 1994). All secondary antibodies were used at 1:2000 Manders’ coefficients were calculated on deconvolved regions of interests of image stacks containing a whole parasite using the scikit-image python package (Walt et al. 2014). Data were analyzed for statistical significance using Kruskal–Wallis comparison test. Chromatic calibration of the microscope was performed prior to imaging experiments.

### Quantitative PCR

Parasite cultures were tightly synchronized using Percoll gradient purification as previously described. Approximately 50 mL of late trophozoite-stage parasites (∼36 h post-invasion) were pelleted and lysed with saponin. Total RNA was extracted from the parasite pellet using the Illustra RNAspin Mini Kit (GE Healthcare, 25-0500-71) according to the manufacturer’s instructions. Purified RNA was reverse-transcribed into complementary DNA (cDNA) using M-MLV Reverse Transcriptase (Promega, M1705).

Real-time quantitative PCR (qPCR) was performed using SsoAdvanced Universal SYBR Green Supermix (Bio Rad, #1725272). Optimal annealing temperatures and primer efficiencies were determined for each primer pair. Reactions were run at 95°C for 5 min, followed by 40 cycles of 95°C for 15 s, the primer-specific annealing temperature for 15 s, and 72°C for 15 s. Melting curve analysis was carried out from 65°C to 95°C at a ramp rate of 0.5°C per 0.5 s. Each qRT PCR reaction was performed in triplicate for each biological replicate, and expression levels were normalized to two reference genes, 60S and tRNA. Data analysis was conducted using the Do my qPCR Calculation tool (Tournayre et al. 2019).

## Author Contributions

F.L. performed experimental work, interpreted the results, and wrote the manuscript. D.S. and J.P.F.M performed experimental work and interpreted the results. P.G and C.R.M designed experiments, interpreted the results and edited the manuscript. D.R. conceived the study, designed experiments, interpreted the results, and wrote the manuscript.

## Acknowledgements

We would like to thank Laura Girardet for her help and advice regarding qPCR and Sabrina Absalon for her advice regarding PfSec14-1 localization. We would also like to thank Tobias Spielman for the SLI and KS plasmids, Jeffrey Dvorin for the pKD and the PfPPP8 plasmids and for antibodies and Alan Cowman for antibodies. We also thank Jacobus Pharmaceuticals for WR99210. The following reagents were obtained through MR4 as part of the BEI Resources, National Institute of Allergy and Infectious Diseases, National Institutes of Health, USA: Polyclonal Anti-*Plasmodium falciparum* PfERD2 (antiserum, Rabbit) and DSM1 (MRA-1161). We would also like to acknowledge the Canadian Blood Services for providing human erythrocytes. UCSF ChimeraX, developed by the Resource for Biocomputing, Visualization, and Informatics at the University of California, San Francisco, with support from National Institutes of Health R01-GM129325 and the Office of Cyber Infrastructure and Computational Biology, National Institute of Allergy and Infectious Diseases. This study was funded through a Canadian Institutes for Health Research project grant (406675) to DR. F.L. received a PhD scholarship from the Fonds de la recherche du Québec-Santé (FRQ-S). DR was a Fonds de la Recherche du Québec-Santé Senior fellow. Work in the McMaster lab was supported by a National Sciences and Engineering Council of Canada Discovery grant (RGPIN 06491). Research in the Griac laboratory is supported by the Scientific Grant Agency of the Ministry of Education, Research, Development, and Youth of the Slovak Republic and the Slovak Academy of Sciences grants VEGA 2/0047/23 and 2/0075/26. DS was supported by Stefan Schwarz Support Fund of the Slovak Academy of Sciences 2024/OV2/011.

## Ethics Statement

Study approved by the Canadian Blood Services (CBS) research ethics board, project number 2023.030 and by the CHU de Québec IRB, project number 2015–2230, B14-12-2230, SIRUL 104595. Written consent was obtained by the CBS for all study participants. Participants were informed about the study before providing consent. All experiments were performed in accordance with relevant guidelines and regulations.

## Data Availability Statement

The data that supports the findings of this study are available in the supplementary material of this article.

## Copyright Statement

No copyright permissions were needed to be taken.

## Tables and figures

## Supporting information

Table S1. List of primers used

**Fig. S1.**
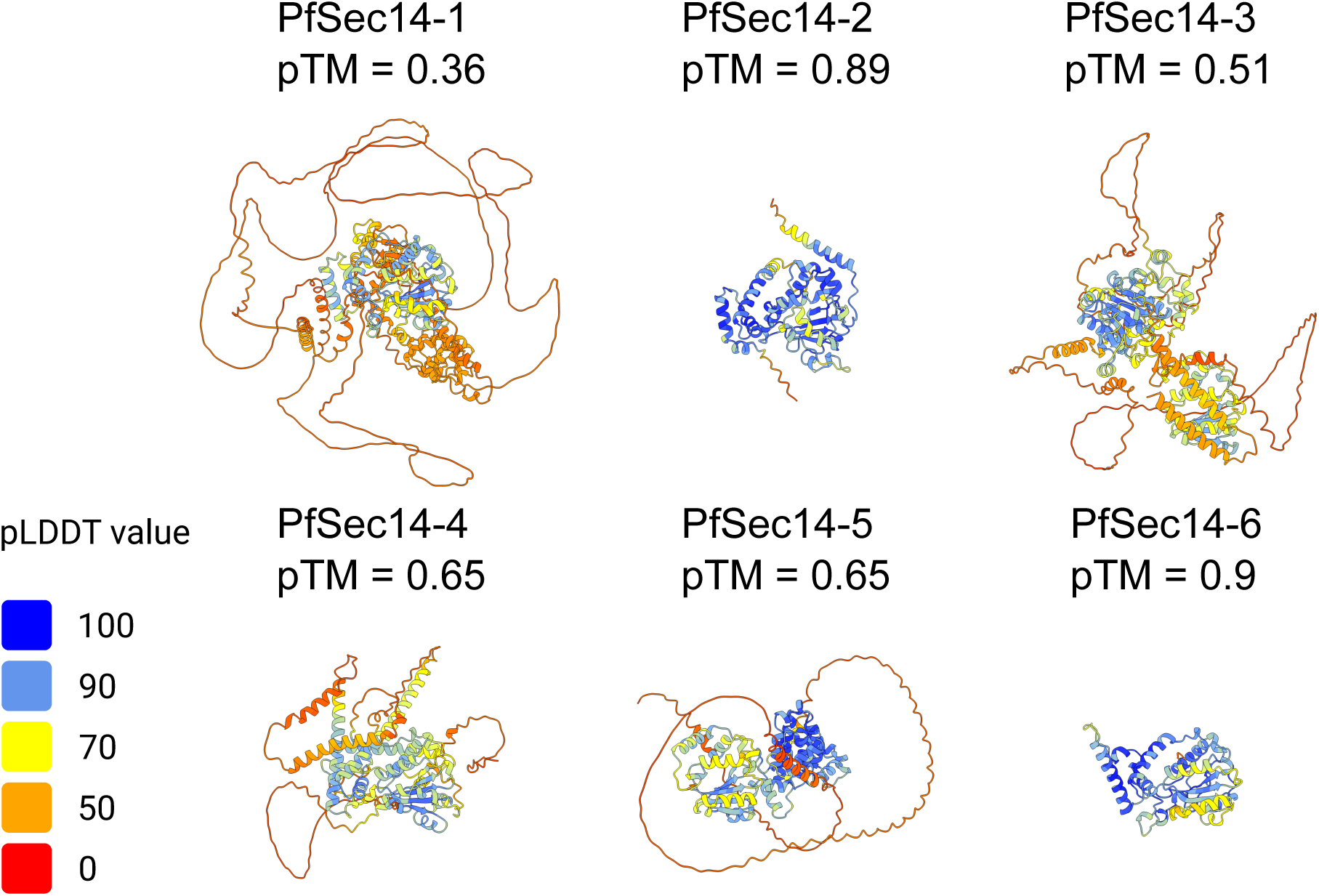
AlphaFold3 structure predictions of PfSec14-1, PfSec14-2, PfSec14-3, PfSec14-4, PfSec14-5 and PfSec14-6. 3D structures are colour coded with predicted local-distance difference test (pLDDT) value.

**Fig. S2.**
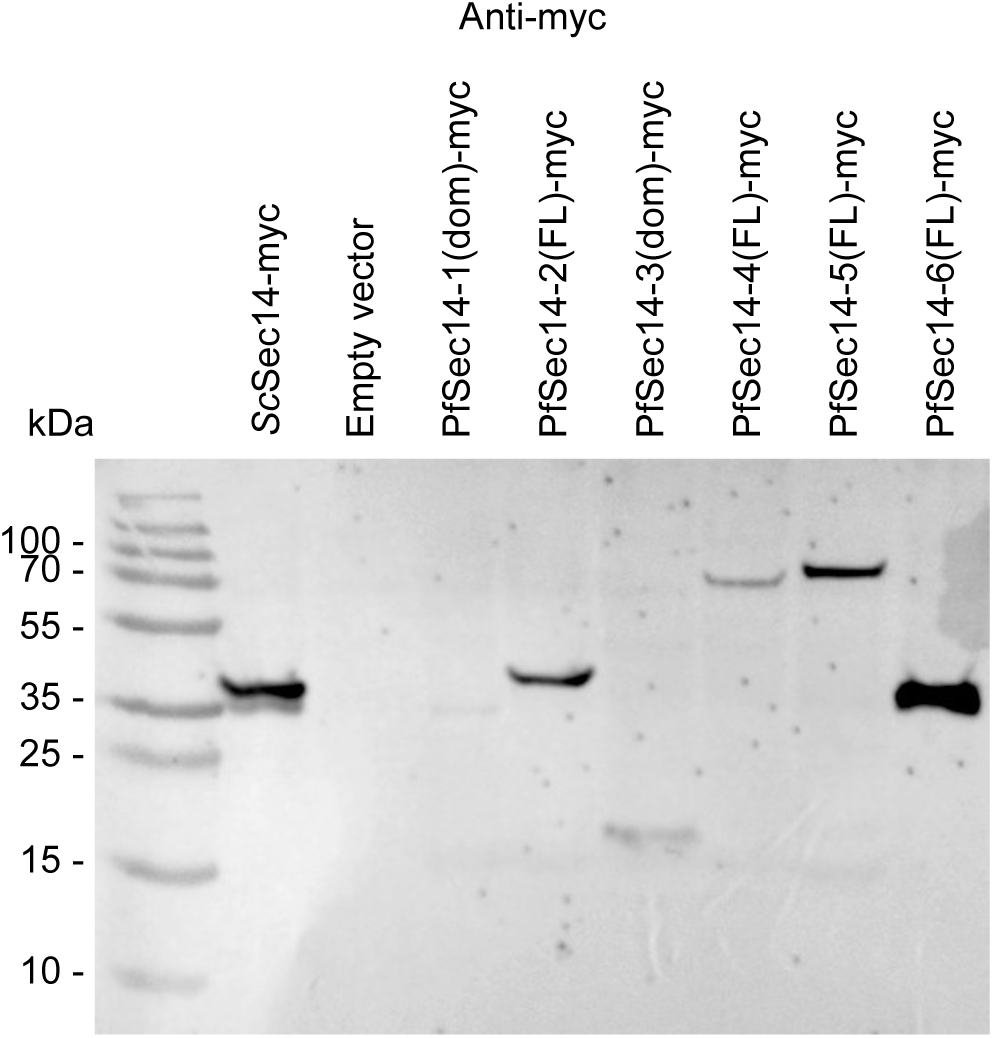
Western blot using an anti-myc antibody showing the expression of the PfSec14 proteins in *S. cerevisiae* at the expected size.

**Fig. S3.**
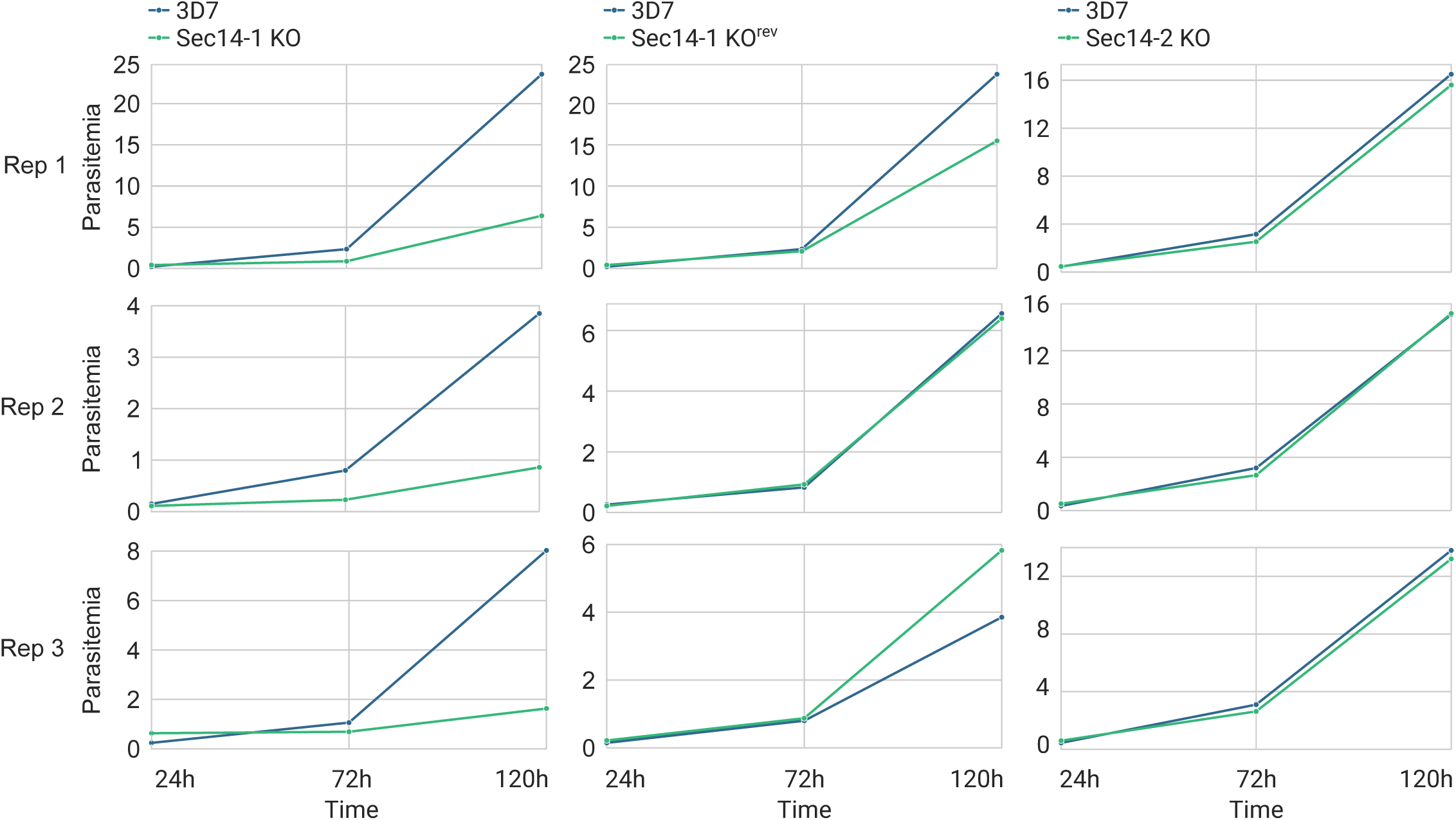
All replicates of flow cytometry growth curves for PfSec14-1 KO and PfSec14-2 KO

**Fig. S4.**
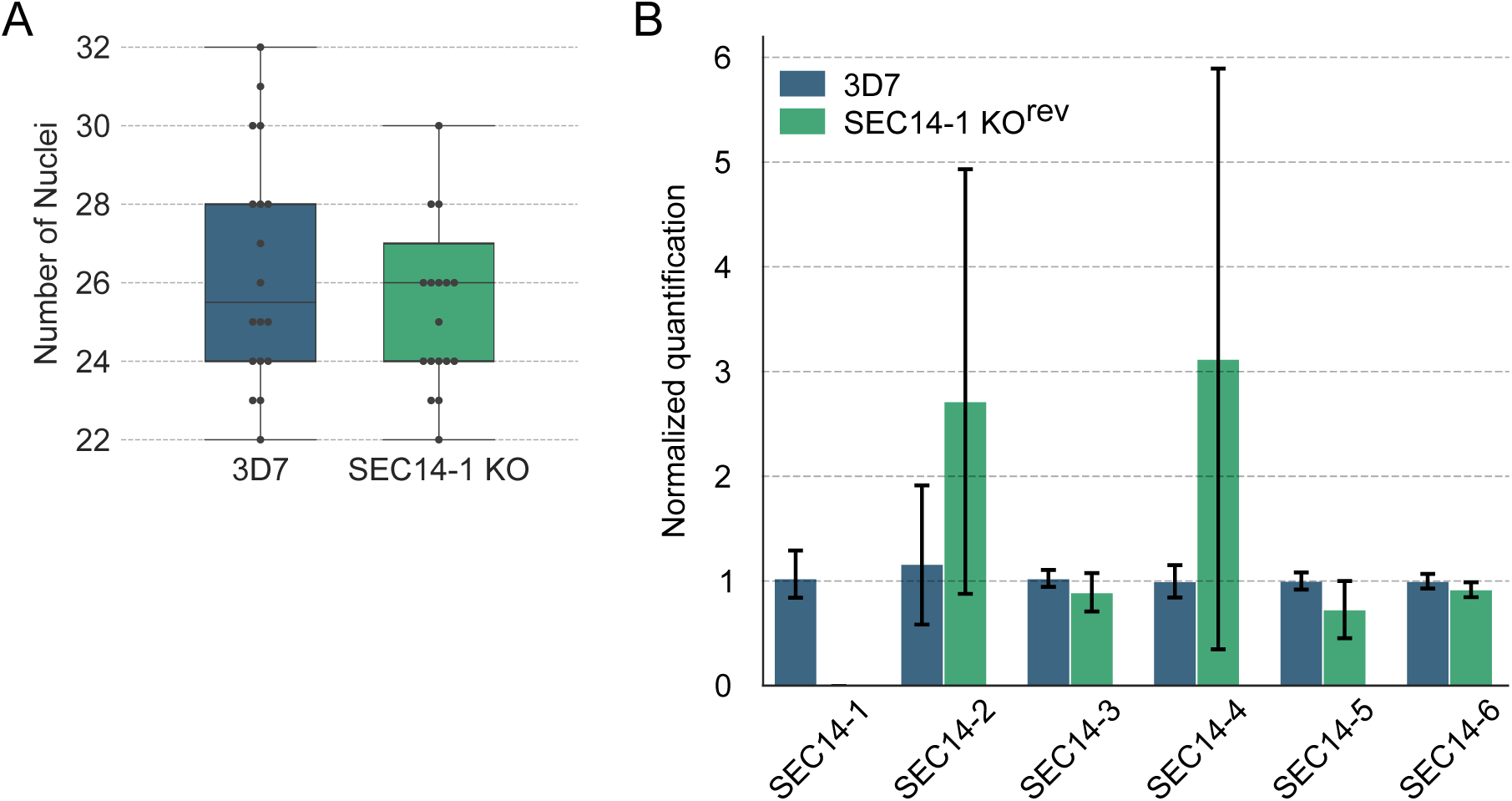
PfSec14-1 TGD strain characterization. (A) Number of merozoites per ML10-arrested schizont in the PfSec14-1 TGD line compared with 3D7. (B) qPCR analysis showing the expression levels of each *PfSEC14* gene in the PfSEC14-1^rev^ line.

**Fig. S5.**
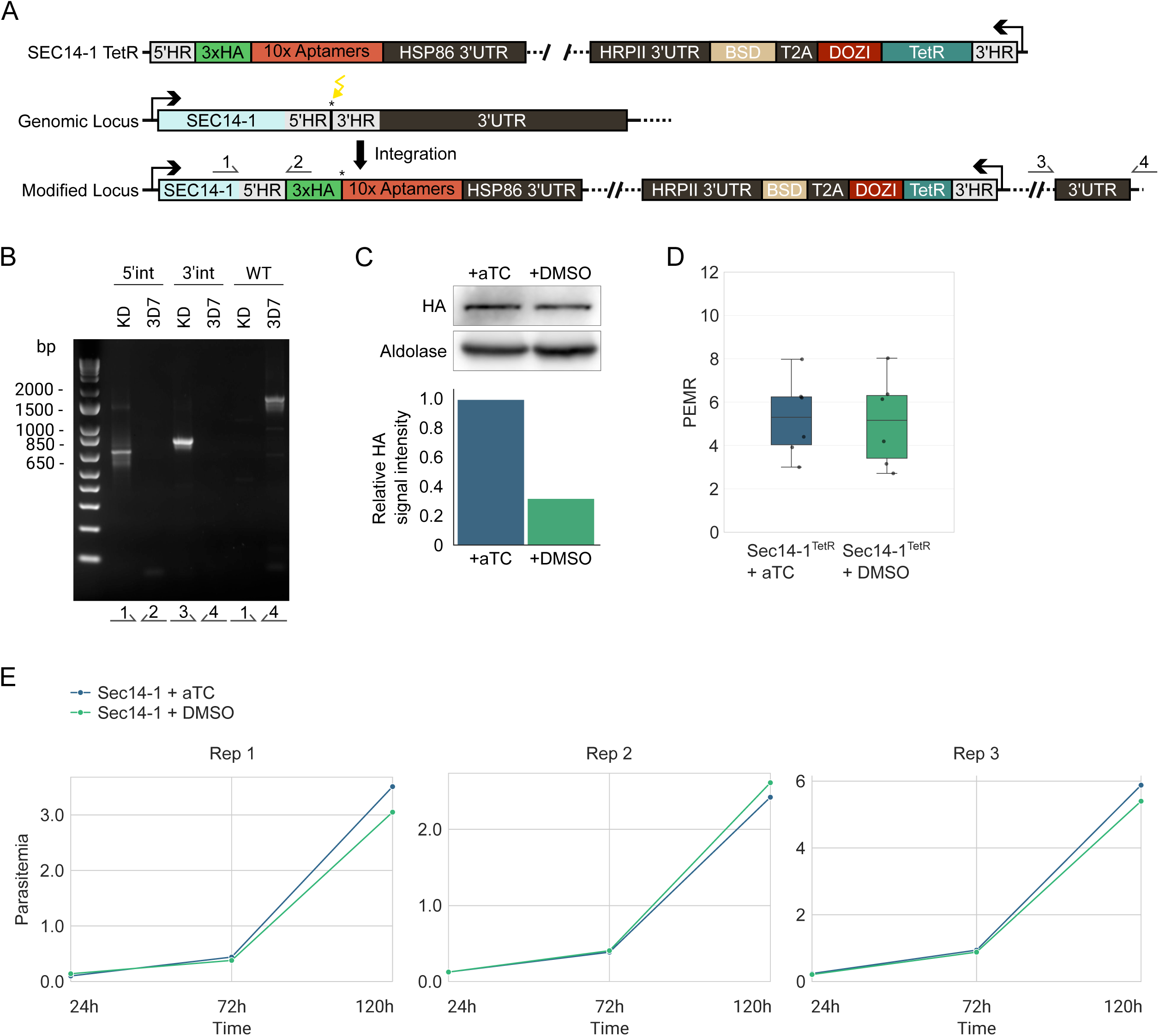
Generation of the PfSec14-1^TetR^ line and induction of PfSEC14-1 knockdown. (A) Illustration of the strategy used to generate the line. Numbered arrows indicate the positions of primers used to verify integration. (B) PCR-based validation of tagging vector integration at the PfSEC14-1 locus performed on parasite genomic DNA from a clonal line. (C) Western blot of protein extracts from trophozoite-stage parasites cultivated for 144 h (three full intraerythrocytic cycles) with or without anhydrotetracycline (aTC). Relative HA signal intensity shows that PfSEC14-1 expression is decreased by approximately 70% in the absence of aTC. (D) Box plot showing growth rate as parasite erythrocyte multiplication rate (PEMR). Each condition was analyzed from three biological replicates. PfSec14-1^TetR^ mean PEMR = 5.10 ± 2.08 (+ aTC) and 5.29 ± 1.84 (+ DMSO). No statistically significant difference was observed between non induced (+ aTC) and non induced (+ DMSO) conditions (Mann–Whitney U test) (E) Individual replicates of flow cytometry–based growth curves of the PfSec14-1^TetR^ line. HR: homology region. HA: hemaglutinnin. UTR: untranslated region. BSD: blasticidin resistance. T2A: 2A peptide. TetR: tetracyclin repressor. aTC: anhydrotetracyclin. DMSO: dimethyl sulfoxide.

**Fig. S6.**
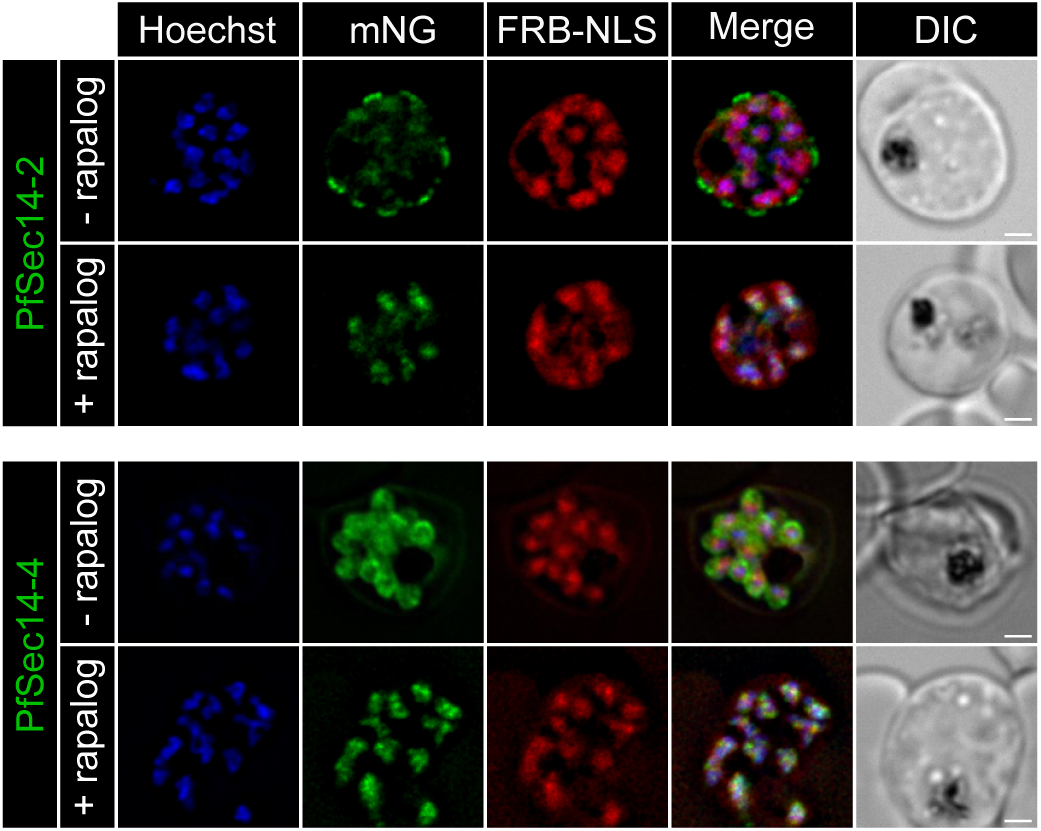
Live cell imaging of PfSec14-2-2xFKBP-mNG + FRB-3xNLS-mCh mislocalizer (mis) and PfSec14-4-2xFKBP-mNG + mis parasites cultivated with or without rapalog, showing efficient rapalog-dependent mislocalization of both PfSec14-2-2xFKBP-mNG and PfSec14-4-2xFKBP-mNG to the nucleus. Scale bar represents 5 µm. Blue: Hoechst-stained nucleus.

**Fig. S7.**
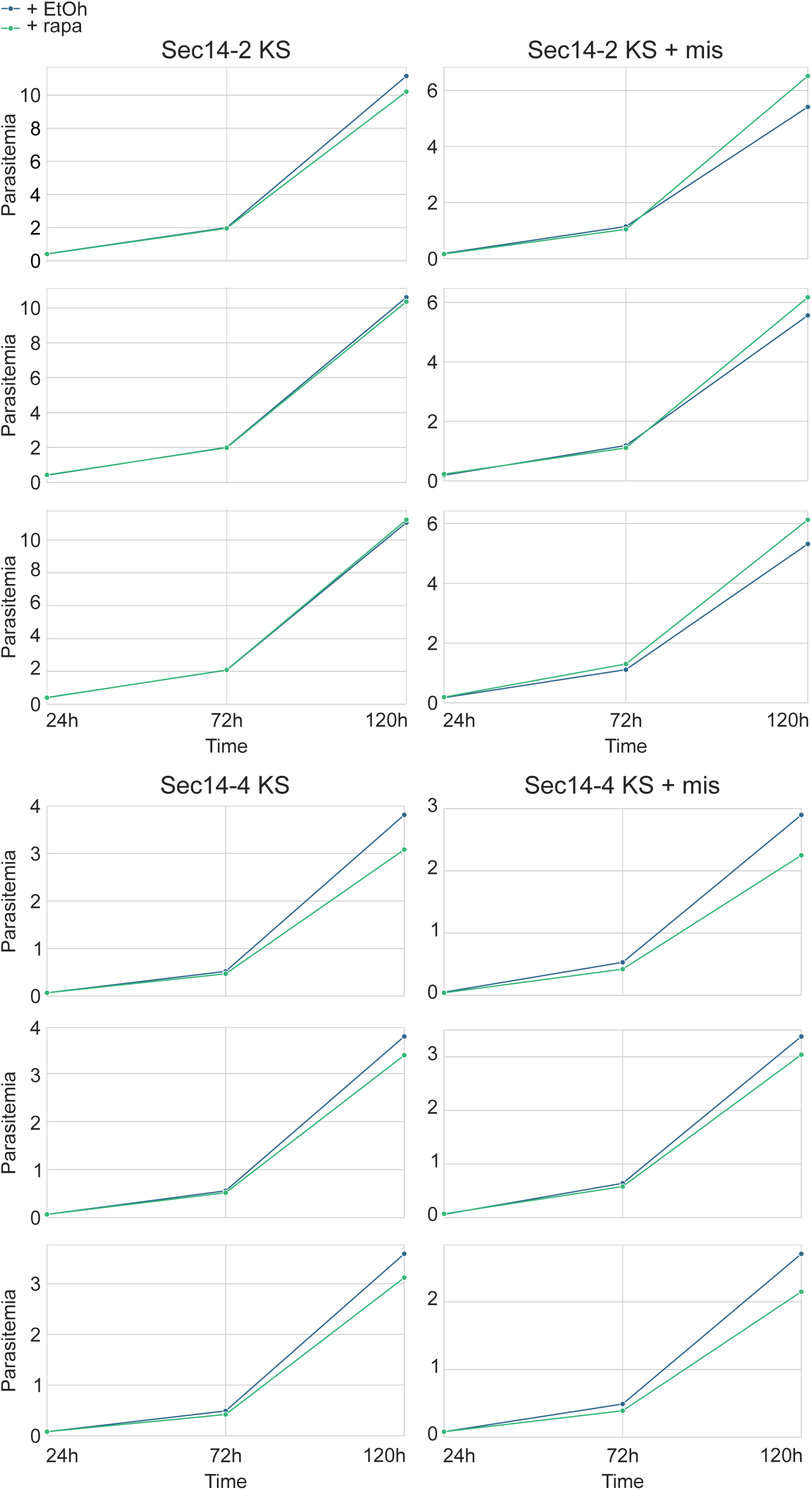
All replicates of flow cytometry growth curves for PfSec14-2 KS and PfSec14-4 KS Appendix 1. Uncropped western blots of protein extracts from ring, trophozoite, and schizont stages of PfSec14-1^TetR^, PfSec14-1 KS, PfSec14-2 KS, and PfSec14-4 KS lines.

