## Supplementary figures and images for "Characterization of Sec14 domain–containing proteins in the malaria parasite *Plasmodium falciparum*"

### Appendix

anti-mNG

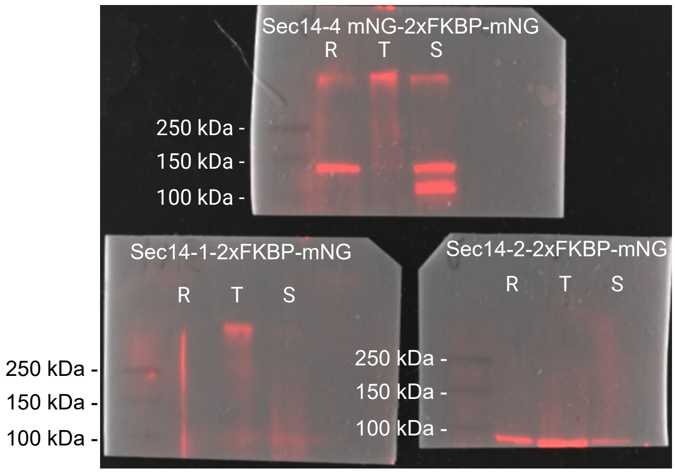


anti-HA

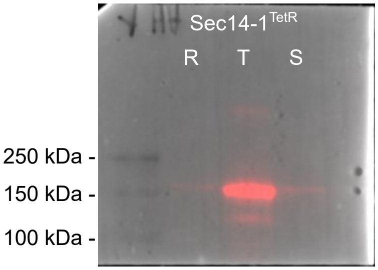


anti-HSP70

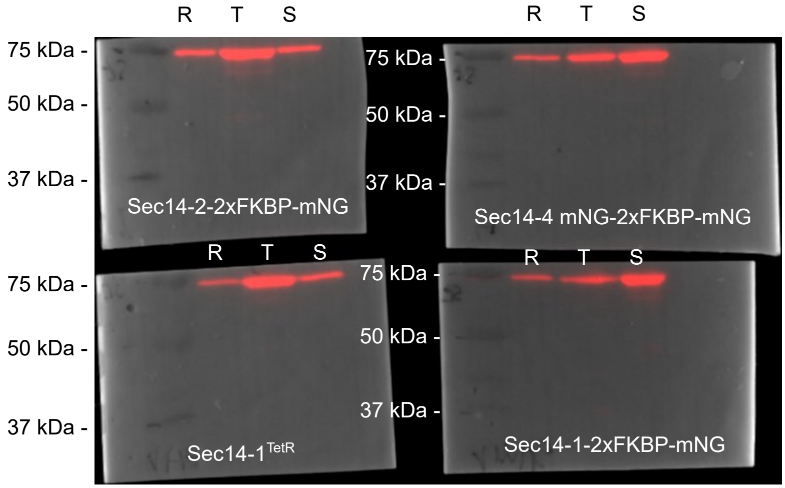
